# Flattened brain hierarchy and increased EEG complexity during unusual bodily experiences and out-of-body experiences

**DOI:** 10.64898/2026.07.30.741513

**Authors:** Teresa Campillo-Ferrer, Tomas Berjaga-Buisan, Antonella Iadarola, Jakub Vohryzek, Ramona Cordani, Marco Veneruso, Yonatan Sanz Perl, Lino Nobili, Robert Oostenveld, Morten L. Kringelbach, Gustavo Deco

## Abstract

Experiences in which the sense of the body becomes distorted offer a unique window into how the brain constructs the bodily self. Despite their clinical relevance in conditions such as depersonalization and body-image disorders, large-scale brain dynamics underlying unusual bodily experiences (UBEs) remain poorly understood, and no prior study has examined them systematically across wakefulness and sleep. Here, we analyzed a high-density EEG dataset including 36 UBE episodes, such as floating sensations, distorted body boundaries, and out-of-body experiences, recorded across rapid eye-movement (REM) sleep, light sleep, sleep arousals, and wakefulness during meditation (N=20). We quantified the temporal irreversibility of neural dynamics, a marker of cortical hierarchical organization, and complemented these analyses with whole-brain signal complexity measures following a within-subject design. UBEs were consistently associated with a global reduction in neural irreversibility and cortical hierarchical differentiation relative to non-UBEs and wakefulness, along with a concurrent increase in whole-brain complexity to levels comparable to wakefulness. Together, these findings suggest that UBEs reflect a neural state combining high informational richness with a transient flattening of cortical hierarchy, a profile that resembles that observed in psychedelic states.

**Key points:**

- UBEs are associated with a global flattening of brain hierarchy measured by EEG.
- UBEs are associated with increased global EEG complexity, resembling the psychedelic state.

## Introduction

Body experience is a fundamental dimension of consciousness and a central focus of neuroscience research. Alterations in this process can lead to states in which body perception is no longer consistent with ordinary wakefulness. This is the case of unusual bodily experiences (UBEs), which involve illusory bodily perceptions that do not align with our everyday waking experience (Campillo-Ferrer *et al*., 2026; Cheyne, 2003; Cheyne *et al*., 1999). For example, individuals during UBEs may report seeing the room with their eyes closed or feeling the bed beneath them when lying down, while simultaneously perceiving themselves as if floating, lacking their body boundaries, or being located outside of their physical body—the latter also known as out-of-body experiences (Alcaraz-Sánchez *et al*., 2022; Blanke *et al*., 2000; Cheyne & Girard, 2009; Levitan *et al*., 1999; Rabeyron & Caussie, 2016).

Although UBEs are frequently reported during sleep and altered states of consciousness, their underlying neural mechanisms remain poorly understood. Early neuroimaging studies relied primarily on anecdotal reports from sleep or relaxation (Levitan *et al*., 1999; Smith & Messier, 2014; Tart, 1967, 1968; Terzaghi *et al*., 2012), or observations from direct brain stimulation in clinical populations (Blanke *et al*., 2000, 2002, 2004; De Ridder *et al*., 2007). While these studies provided important initial insights, they were limited in both reproducibility and scope. Recent methodological advances have allowed UBEs to be investigated systematically under controlled conditions. For instance, eye movement tracking has been used to identify spontaneous experiences during sleep (Herrero *et al*., 2025; Mainieri *et al*., 2021; Weiler *et al*., 2025). In parallel, a wide variety of induction methods have been used to simulate UBEs or facilitate their appearance under controlled conditions, including sleep interruption techniques, hypnosis, meditation, psychedelics, sensory limitation, multisensory conflict, or virtual reality (Dor-Ziderman *et al*., 2016; Ionta *et al*., 2011; Lenggenhager *et al*., 2011; Martial *et al*., 2023; McCreery & Claridge, 1996; Tagliazucchi *et al*., 2016; Takeuchi *et al*., 1992, 2002; Tressoldi *et al*., 2015; Zeev-Wolf *et al*., 2017).

Building on these methodological advances, the neural correlates of UBEs have been investigated using electroencephalography (EEG), magnetoencephalography (MEG) and functional magnetic resonance imaging (fMRI). Studies using fMRI have consistently implicated regions involved in multisensory integration and body self-perception, such as the temporo-parietal junction and insular cortex (Blanke & Arzy, 2005; Ionta *et al*., 2011). In contrast, electrophysiological studies show more heterogeneous findings: some report reduced alpha power (Lenggenhager *et al*., 2011; Martial *et al*., 2023), while others have found increases in alpha, delta or theta activity (Berkovich-Ohana *et al*., 2013; Martial *et al*., 2023; Zeev-Wolf *et al*., 2017), and decreases in beta and gamma activity in similar or overlapping regions (Dor-Ziderman *et al*., 2016; Tressoldi *et al*., 2015; Zeev-Wolf *et al*., 2017).

While these findings may appear contradictory at first, they could also reflect a common underlying tendency: a shift toward EEG deactivation that could manifest as increased power in lower frequencies (like delta and theta) and/or reduced power in higher frequencies (like beta and gamma), a spectral profile that resembles neural dynamics typically observed at sleep onset (Marzano *et al*., 2013; Siclari *et al*., 2014). This raises the possibility that spectral markers attributed to UBEs may not represent unique signatures of the phenomenon, but instead reflect mixed wake-sleep electrophysiology that is not well captured by conventional sleep-staging criteria. Consistent with this view, several studies have observed UBEs arising during hybrid EEG states combining elements of both wakefulness and sleep (Campillo-Ferrer *et al*., 2026; Mainieri *et al*., 2021; Takeuchi *et al*., 1992, 2002; Terzaghi *et al*., 2012).

Taken together, these findings call for methodological approaches that investigate the brain’s global organization during UBEs, rather than focusing solely on spectral changes. Promising steps in this direction include reports of altered functional and effective connectivity in the temporo-parietal junction and insular cortex during UBEs reported in psychedelic states (Stoliker *et al*., 2025; Tagliazucchi *et al*., 2016), or reduced theta-band complexity and increased beta-band connectivity during UBEs simulated through virtual reality (Martial *et al*., 2023). However, these findings need replication in larger and more diverse populations to determine whether they generalize beyond these paradigms. Importantly, none of these studies have examined global patterns across both wakefulness and sleep, limiting our understanding of the overall brain dynamics associated with UBEs.

To address these gaps, we adopted a non-equilibrium perspective on large-scale brain dynamics, conceptualizing neural activity as an asymmetric system evolving far from equilibrium (Kringelbach *et al*., 2024). This framework is grounded in the second law of thermodynamics, which states that a system cannot naturally transition from disorder to order. For example, a broken glass cannot spontaneously reassemble, which makes this process irreversible. Applied to brain signals, this principle implies a preferred direction of information flow, known as the “arrow of time” (Eddington, 1928; Feng & Crooks, 2008; Seif *et al*., 2021), which can be used to quantify the level of irreversibility and hierarchical organization of brain dynamics. This approach has been successfully applied to discriminate and mechanistically characterize different brain states across multiple imagining modalities, consistently revealing increased irreversibility during conscious states like wakefulness and dreaming, in contrast to deep sleep, anaesthesia, and multiple neurological conditions (Berjaga-Buisan *et al*., 2025; Camassa *et al*., 2024; Cruzat *et al*., 2023; Deco *et al*., 2022, 2023; G-Guzmán *et al*., 2023; Idesis *et al*., 2024; Sanz Perl *et al*., 2021).

In this study, we used the INSIDEOUT approach developed by Deco *et al*., (2022) to quantify the irreversibility of high-density EEG signals obtained from a previously published dataset of 20 healthy individuals, including 36 UBE episodes reported across wakefulness and sleep. This approach provides a model-free method for estimating neural irreversibility and hierarchical organization through the quantification of asymmetries in brain signals, specifically by assessing whether time-forward and time-reversed trajectories are distinguishable from each other (Deco *et al*., 2022). We hypothesized that UBEs would be associated with distinct changes in the hierarchical organization of brain activity, potentially revealing markers that distinguish them from ordinary states of consciousness like wakefulness or sleep. To provide a broader perspective on the EEG correlates of UBEs, we complemented irreversibility and hierarchy analyses with Lempel-Ziv (LZ) complexity, a marker of EEG diversity (Aamodt *et al*., 2022; Schartner, Pigorini, *et al*., 2017), and revisited spectral analyses previously performed on the same dataset (see Campillo-Ferrer *et al*., 2026). This allowed us to directly compare approaches and integrate findings across different metrics.

## Materials & methods

### Dataset

The dataset analysed in the present study was originally collected at the Donders Centre for Cognitive Neuroimaging (Radboud University Nijmegen, the Netherlands) and is publicly available at: https://doi.org/10.34973/50bx-st45. It comprises high-density EEG (64 electrodes) and EOG, EMG, and ECG recordings from 35 healthy participants who underwent a single-session UBE induction procedure in a controlled sleep laboratory setting. Here, we analysed signals corresponding to 36 UBEs reported by 20 participants (mean age = 25.14 ± 4.59 years; 15 females, 5 males), as well as signals corresponding to wakefulness, meditation, and sleep from the same participants. The original study received approval from the local medical ethical committee in Nijmegen, the Netherlands (Imaging Human Cognition, NL45659.091.14), under the blanket approval of the Donders Centre for Cognitive Neuroimaging (Donders Institute, the Netherlands). Prior to the experiments, all participants had experienced at least one lucid dream (i.e., dream in which the individual is aware of being in a dream state) or had prior experience with meditation practices (e.g., participation in a university mindfulness course). All participants provided written informed consent prior to participation. Detailed sample characteristics and methodology can be found in the original study (Campillo-Ferrer *et al*., 2026); here we only provide a short summary:

#### Experimental procedures

In the original study, UBEs were defined to participants during an intake session (1-7 days before the experimental session) as “sensations you would not usually experience during normal wakefulness, such as having strong bodily vibrations, floating, or feeling as if you were going outside of your physical body”. Participants could experience them while dreaming or while still being aware of their physical surroundings (e.g., during meditation). During the experimental session, induction procedures were carried out in the following order: (1) mild sleep deprivation (∼5 hours of sleep in the preceding night), (2) semi-guided meditation practice (∼1 hour, starting at 8:30 a.m.) combining square breathing, body scan meditation, the “corpse pose” position used in Yoga Nidrâ, (3) morning nap (∼3 hours), during which participants were asked to continue meditating and retain consciousness while falling asleep, (4) sensory stimulation with light cues presented across wakefulness and sleep (for induction method, see Campillo-Ferrer *et al*., 2025). To time-mark UBEs in the EEG, participants were asked to perform a left-right-left-right (LRLR) eye movement as soon as they experienced an UBE, a signal commonly used in lucid dreaming studies (Demirel *et al*., 2025; LaBerge *et al*., 1981) and more anecdotally to localize UBEs in the EEG (Herrero *et al*., 2025; Mainieri *et al*., 2021; Weiler *et al*., 2025). In parallel, EEG, EOG and EMG signals were continuously monitored in real time.

#### EEG recordings

In the original study, EEG recordings were obtained using 64 active scalp electrodes positioned according to the international 10-10 system (actiCAP, BrainProducts GmbH), referenced to FCz, and sampled at 500 Hz. Additional passive electrodes (one bipolar EOG, two submental EMGs, and one bipolar ECG) were used for sleep staging and auxiliary physiological recordings (BrainAmp ExG, BrainProducts GmbH).

#### Phenomenology

In the original study, participants provided 1-2 minute open experience reports short after performing LRLR eye signals to describe their UBEs, and were then allowed to continue sleeping. At the end of each session, interviews inspired by the micro-phenomenological technique were conducted (Petitmengin, 2006; Petitmengin *et al*., 2019; Varela, 1996) and partly analysed together with the participant. This approach allowed to (1) verify consistency between LRLR eye signals, open experience reports, and interview data; (2) reconstruct the temporal dynamics of each UBE; (3) characterize the phenomenological features of each UBE (Valenzuela-Moguillansky & Vásquez-Rosati, 2019). The final dataset includes UBEs with one or more of the following elements: (1) vestibular-motor experiences; (2) tactile sensations; (3) distortion of body boundaries; (4) lack of bodily sensations and/or position; (5) out-of-body and/or elevated self-location experiences; (6) sleep paralysis, tension and/or immobility experiences.

#### Sleep scoring

In the original study, sleep staging was performed by trained experts in accordance with the American Academy of Sleep Medicine guidelines (AASM manual, 2020) and conducted in two iterative steps: first using 30-second epochs, and then using 5-second mini-epochs. The final dataset includes UBEs reported in four different stages: (1) REM sleep, (2) stages N1 and N2 of non-REM sleep (light sleep), (3) wakefulness during meditation, and (4) wakefulness occurring within 60 seconds after arousals from light sleep (sleep arousals).

### EEG data and preprocessing

Preprocessing for the present analyses followed a procedure similar to the original study, using the FieldTrip toolbox (Oostenveld *et al*., 2011) in MATLAB (Version R2024a, MathWorks Inc.). Briefly, ≥ 400 seconds of eyes-closed wakefulness and ≥ 400 seconds of the conscious state in which UBEs were reported (e.g., REM sleep) were selected, including both UBE and non-UBE episodes. Muscular artifacts and bad channels were identified by visual inspection, EEG data were bandpass filtered (1-45 Hz), bad channels were interpolated (for neighbour structure, see Campillo-Ferrer *et al*., 2026) and remaining windows containing muscular artifacts were rejected. Independent component analysis (ICA) was then applied to remove eye and cardiac artifacts, and data were re-referenced to the mean of both mastoids using TP9 and TP10. Once EEG signals were clean, the data were segmented into three conditions for each participant:

1. <u>UBE condition</u>: 5-second segments centred around each LRLR eye signal (up to 4 per participant). Depending on the phenomenological reports, these segments could include the 5 seconds before the eye signal, after the eye signal, or be split between both (e.g., 2.5 seconds before and 2.5 seconds after). In all cases, 1 second immediately before and after eye signals were excluded to avoid activity related to performing the signal.
2. <u>Wakefulness condition</u>: one 60 second segment of eyes-closed wakefulness recorded at the session onset, before the meditation practice.
3. <u>Non-UBE condition</u>: one 60 second segment of the same sleep or wake state as the UBE, where the segment closest in time to the UBE and free of fragmentation and UBE reports was selected. For UBEs reported after arousals from light sleep, light sleep was selected as the non-UBE condition.

In the present study, data from all conditions were further divided into 1-second non-overlapping sliding windows to capture detailed brain dynamics, resulting in: 177 clean windows for the UBE condition, 1151 windows for the wakefulness condition, and 1365 windows for the non-UBE condition. The signal was additionally mean-centered and linearly detrended to remove residual offsets within each window. To apply the INSIDEOUT framework, the EEG channel dimensionality was reduced using principal component analysis (using the *pca* function in MATLAB) and retaining only the top components that together explained 90% of the channel variance. Raincloud plots were generated using the *daviolinplot* function in MATLAB (Karvelis, 2025) and topographic maps using the FieldTrip toolbox.

### Determining the levels of irreversibility and hierarchy

The level of temporal non-reversibility (i.e., irreversibility), and thus non-equilibrium, is quantified for each 1-second window and EEG channel by detecting the arrow of time through asymmetries in causal interactions between the original and the time-reversed signals, as described in previous studies (INSIDEOUT framework, Camassa *et al*., 2024; Cruzat *et al*., 2023; Deco *et al*., 2022).

To illustrate the methodology, consider the bivariate case between two time series *x*(*t*) and *y*(*t*). The forward time-shifted correlation is defined as:

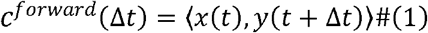

and the reversed time-shifted correlation, obtained after inverting the temporal order of both series, as:

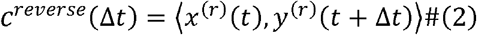

Therefore, the pairwise irreversibility index, capturing degree of temporal asymmetry (i.e., the arrow of time), is given by the absolute difference between the causal relationship between the two time-shifted correlations at a given shift:

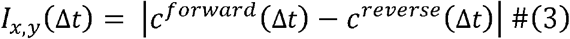

For the multidimensional case, the irreversibility is quantified by constructing forward and reversed matrices of time-shifted correlations, representing the causal dependencies among system variables in the original and time-reversed dynamics:

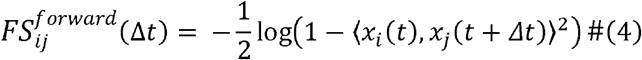

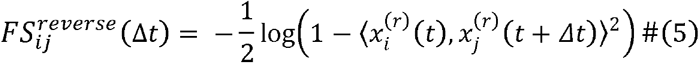

These matrices correspond to the mutual information of time-shifted correlations. The global degree of irreversibility is quantified as the quadratic distance between the forward and reversed matrices at a given time lag *∆t* = *T*, where the optimal time lag T was determined as the point where the averaged autocorrelation (AC) across conditions has sufficiently decayed (AC = 10% of its maximum, corresponding to a 0.064 s time shift):

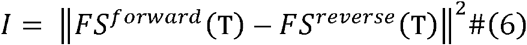

In other words, the level of irreversibility is calculated as the mean value of the elements of the squared-difference matrix:

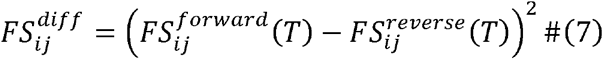

Finally, the hierarchy is quantified as the standard deviation of the elements of the squared-difference matrix, reflecting the heterogeneity of causal asymmetries across brain regions. This measure captures the degree of hierarchical differentiation in the system’s non-equilibrium dynamics.

### Determining the levels of complexity

Lempel-Ziv (LZ) complexity was estimated for each 1-second window and EEG channel using the *kolmogorov* function in MATLAB (Faul, 2025). Each time series was first binarized by converting signal values above the median to “1” and those below the median to “0”. The function then applies the algorithm originally described by Lempel & Ziv (1976) and later adapted for timeseries analysis by Kaspar & Schuster (1987). In brief, the algorithm scans the binarized signal and counts the number of different patterns or sub-sequences present in the signal. Repetitive signals present low LZ complexity values, while more irregular or unpredictable signals present higher values, indicating greater EEG signal diversity (Aamodt *et al*., 2022; Schartner, Pigorini, *et al*., 2017). Global LZ complexity was calculated as the mean complexity value across channels and was used for subsequent statistical comparisons.

### Statistics

Generalized linear mixed models (GLMMs) were used to compare values across experimental conditions, following a within-subject design, and using the *fitglme* function in MATLAB (R2024b, MathWorks Inc.). Random effects for *subject number* and *conscious state* were included to account for (1) the unequal number of trials across subjects and (2) the different states of consciousness in which UBEs were reported. Models were compared iteratively, starting from the simplest structure (i.e., null model including only the intercept) and progressively adding fixed effects (*experimental condition*) and different combinations of random intercepts and slopes (*subject number*, *conscious state*, and/or their interaction). Only effects that significantly improved model fit were retained, as determined by likelihood ratio tests using the *compare* function in MATLAB. The final models included *experimental condition* as a fixed effect and the interaction of *subject number* x *conscious state* as a random intercept. Models were specified using a Gamma distribution with a log link function (for irreversibility and hierarchy) and a Normal distribution with an identity link function (for complexity), and were fitted using the Laplace method. For analyses in which the spectrum was divided into frequency bands, one separate model was fitted per frequency band (six in total). To correct for multiple comparisons, the false discovery rate (FDR) was applied using the *mafdr* function in MATLAB.

## Results

To test our main hypothesis, we characterized the brain’s hierarchical organization of 36 UBEs reported by 20 participants across wakefulness and sleep, using high-density EEG signals and comparing them to resting wakefulness (eyes-closed) and non-UBE episodes from the same participants. To ensure state-matched comparisons, the non-UBE condition was defined as the same state of consciousness in which UBEs were reported, but in the absence of any UBE occurrence. This included REM sleep, light sleep, and wakefulness during meditation (see Methods). In the case of UBEs reported during sleep arousals (all originating from light sleep), a segment of light sleep was selected as the corresponding non-UBE condition.

UBEs were facilitated using a meditation and light stimulation protocol (**Figure 1A**), resulting in a wide variety of vestibular-motor and tactile sensations, episodes with distorted body boundaries or absent bodily awareness, out-of-body experiences, and sleep paralysis episodes (**Figure 1B**). Each experience was time-wise localized in the EEG using a predefined left-right-left-right eye movement sequence performed by participants. These movements were clearly visible in the auxiliary EOG channels and served as objective markers for UBE onset and offset (**Figure 1C**). EEG segments of 5 seconds were extracted around these signals and further validated through interviews inspired by the micro-phenomenological technique (for further details see Campillo-Ferrer *et al*., 2026).

**Figure 1.**
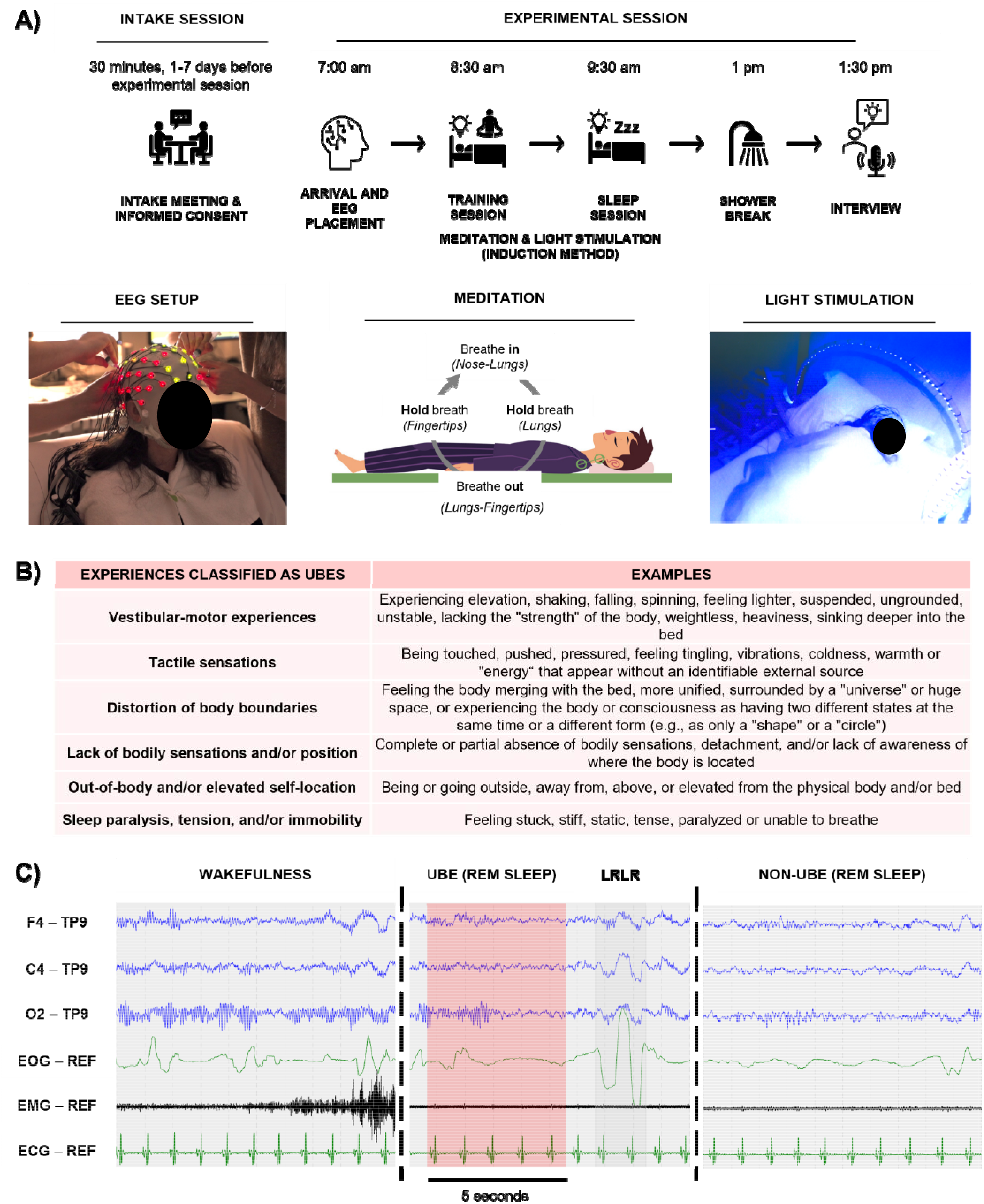
Experimental setup, protocol, and example data. A) We used an existing dataset of n=36 UBEs reported by 20 subjects across different states of consciousness: (1) REM sleep, (2) stages N1 and N2 of non-REM sleep (light sleep), (3) wakefulness during meditation, and (4) wakefulness following arousals from light sleep (sleep arousals). The protocol consisted of a 30-minute intake session conducted days before the experiment to obtain informed consent and information, followed by a 6.5-hour experimental session with live EEG monitoring and an induction protocol combining meditation and light stimulation to facilitate UBEs. Photographs of the EEG setup and light stimulation were taken by Paula Ábalos as part of her audiovisual project “APARICIONES”: (https://apariciones.net/ and https://paulaabalos.com/APARICIONES). The participant consented to the use of these photographs. Illustration of a man lying down to exemplify meditation: © Can Stock Photo Inc. / [sabelskaya]. B) Examples of experiences classified as UBEs. Classification was based on open experience reports (collected right after the experience) and interviews inspired by the micro-phenomenological technique (collected at the end of the session). C) Example of frontal, central and occipital EEG activity, alongside EOG, EMG and ECG signals for three experimental conditions within the same subject: wakefulness (eyes-closed, sleep-deprived 2-3h), UBE (in this case reported during REM sleep), and non-UBE (REM sleep without reported UBEs). Left-right-left-right eye movements (LRLR, dark grey shading) were used as objective markers for UBEs, while 5-second segments corresponding to UBEs (pink shading) were used for all EEG analyses.

Brain irreversibility was estimated using the INSIDEOUT framework developed by Deco *et al*. (2022), which compares the causal relationship between pairwise time series in the forward EEG signal versus an artificially generated reversed version at a given time shift T (**Figure 2**). For the current study, we selected the value of T at which the average autocorrelation across conditions has decayed to 10% of its maximum value. Brain hierarchy was estimated as the variability (standard deviation) of brain irreversibility across different cortical areas. Finally, statistical analyses were performed using generalized linear mixed models (GLMMs), where *subject number* was included as a random effect to account for repeated measures within conditions, and *conscious state* to account for variability across different states of consciousness.

**Figure 2.**
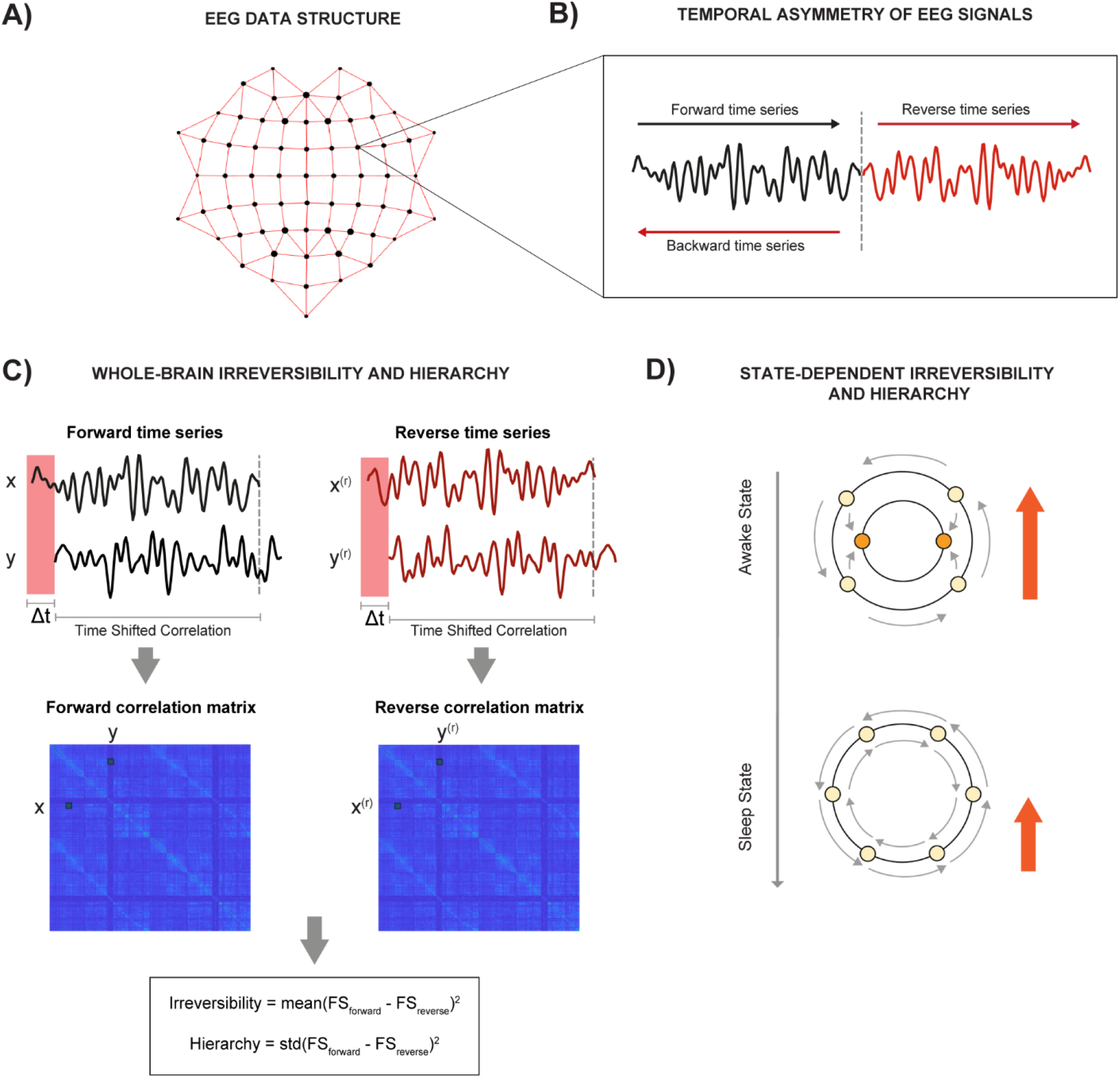
Method for estimating temporal asymmetries in EEG signals. A) EEG activity was recorded using a 64-electrode cap and subsequently projected onto a lower-dimensional space by principal component analysis (PCA), retaining the components that explain 90% of the total channel variance. B) A time-reversed EEG signal is generated by computing the backward (mirror) version of the original time-forward signal. C) Temporal asymmetry is quantified by computing shifted correlations for each pair of time series *x*(*t*) and *y*(*t*) for a given temporal shift *∆t* = *T*. This process is repeated separately for both the forward and time-reversed EEG signals, *x*^*(r)*^(*t*) and *y*^*(r)*^(*t*), resulting in forward and reversed correlation matrices, *FS_forward_* and *FS_reverse_*. Irreversibility and hierarchy are then defined as the mean (irreversibility) and standard deviation (hierarchy) of the elements of the squared difference between these two matrices. D) Awake and conscious states are typically associated with higher irreversibility and hierarchy, reflecting increased temporal asymmetry and the dominance of specific brain regions over global brain activity, whereas deep sleep and unconscious states are commonly associated with lower values, indicating more temporally symmetric dynamics and a more homogeneous contribution of brain regions.

### Broadband EEG irreversibility

Broadband EEG irreversibility (1-45 Hz, **Figure 3A**) was significantly reduced during UBEs compared to non-UBE states (FDR-adjusted p < 0.001). By contrast, no significant differences were found between wakefulness and either the UBE (FDR-adjusted p = 0.14) or non-UBE conditions (FDR-adjusted p = 0.78). A similar pattern was observed for hierarchy measures: values were significantly lower in UBE compared to non-UBE episodes (FDR-adjusted p < 0.001), but wakefulness did not differ from either UBE (FDR-adjusted p = 0.22) or non-UBE states (FDR-adjusted p = 0.71). Topographic maps further showed that the same brain regions (mainly occipital and right-frontocentral) contributed to irreversibility in all conditions, with a gradual tendency toward decreased intensity from wakefulness to non-UBE states and UBE episodes.

**Figure 3.**
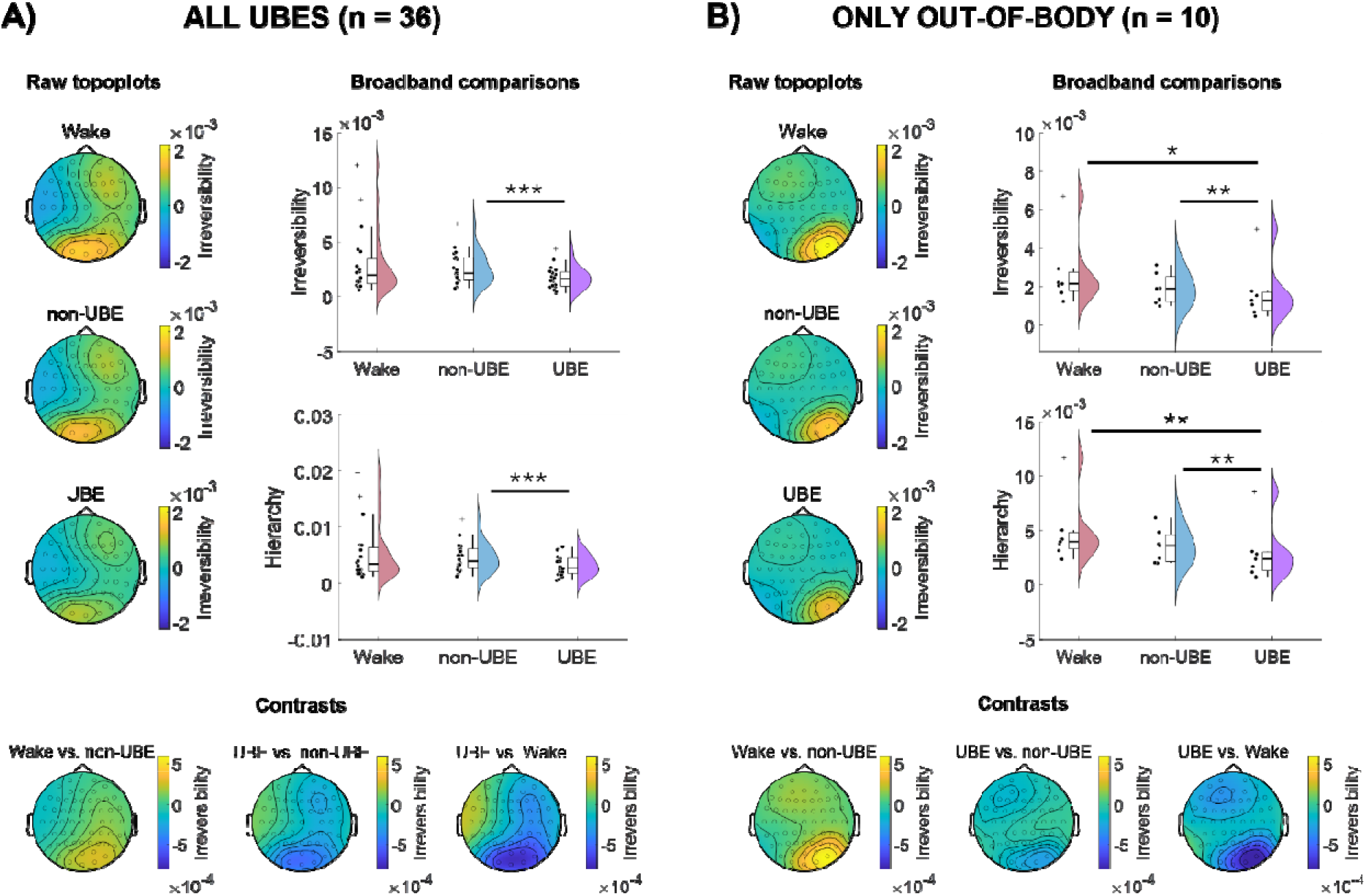
Broadband EEG irreversibility across conditions. A) Full UBE sample (n=36 UBEs, N=20 subjects). B) UBEs involving out-of-body and/or elevated self-location experiences (n=10 UBEs, N=7 subjects). Dots indicate subject-level means for irreversibility (top) and hierarchy (bottom) for visualization. Statistical comparisons with generalized linear mixed models were performed on individual trials and following a within-subject design. Topographic maps show the spatial distribution of irreversibility for individual conditions (left) and contrasts (bottom). Significance of model comparisons: *** p < 0.001, ** p < 0.01, * p < 0.05.

When restricting the analysis to UBEs involving out-of-body experiences and feelings of elevation (n=10 UBEs from N=7 subjects), similar results were obtained (**Figure 3B**). These episodes showed a significant decrease in both irreversibility and hierarchy compared to non-UBEs (FDR-adjusted p < 0.01). This time, both measures were also significantly reduced in UBEs relative to wakefulness (FDR-adjusted p < 0.01 for hierarchy, FDR-adjusted p < 0.05 for irreversibility). As with the full UBE sample, the brain regions sustaining irreversibility remained largely unchanged across conditions, again showing a tendency toward a gradual decrease in intensity from wakefulness to non-UBE states and UBE episodes, this time most evident in right-occipital and left-frontocentral areas.

The results remained consistent when comparing UBEs and non-UBEs across three different settings (sensitivity analyses shown in **Figure 4**): (1) when testing different values of T at which the average autocorrelation (AC) has decayed to 0%, 10%, 20%, 30%, 40%, and 50% of its maximum (FDR-adjusted p < 0.001 in all cases); (2) when considering only the top 20% of irreversibility values (FDR-adjusted p < 0.001 in all cases); and (3) when splitting the dataset into four subgroups according to the conscious state in which UBEs were reported (FDR-adjusted p < 0.05 in all cases except for the ‘light sleep’ group, where p = 0.054). Differences between UBEs and wakefulness were observed when T was set at AC = 50%, as well as when UBEs were analyzed separately for the ‘REM sleep’, ‘light sleep’, and ‘meditation’ groups (see Supplementary Tables for detailed statistics).

**Figure 4.**
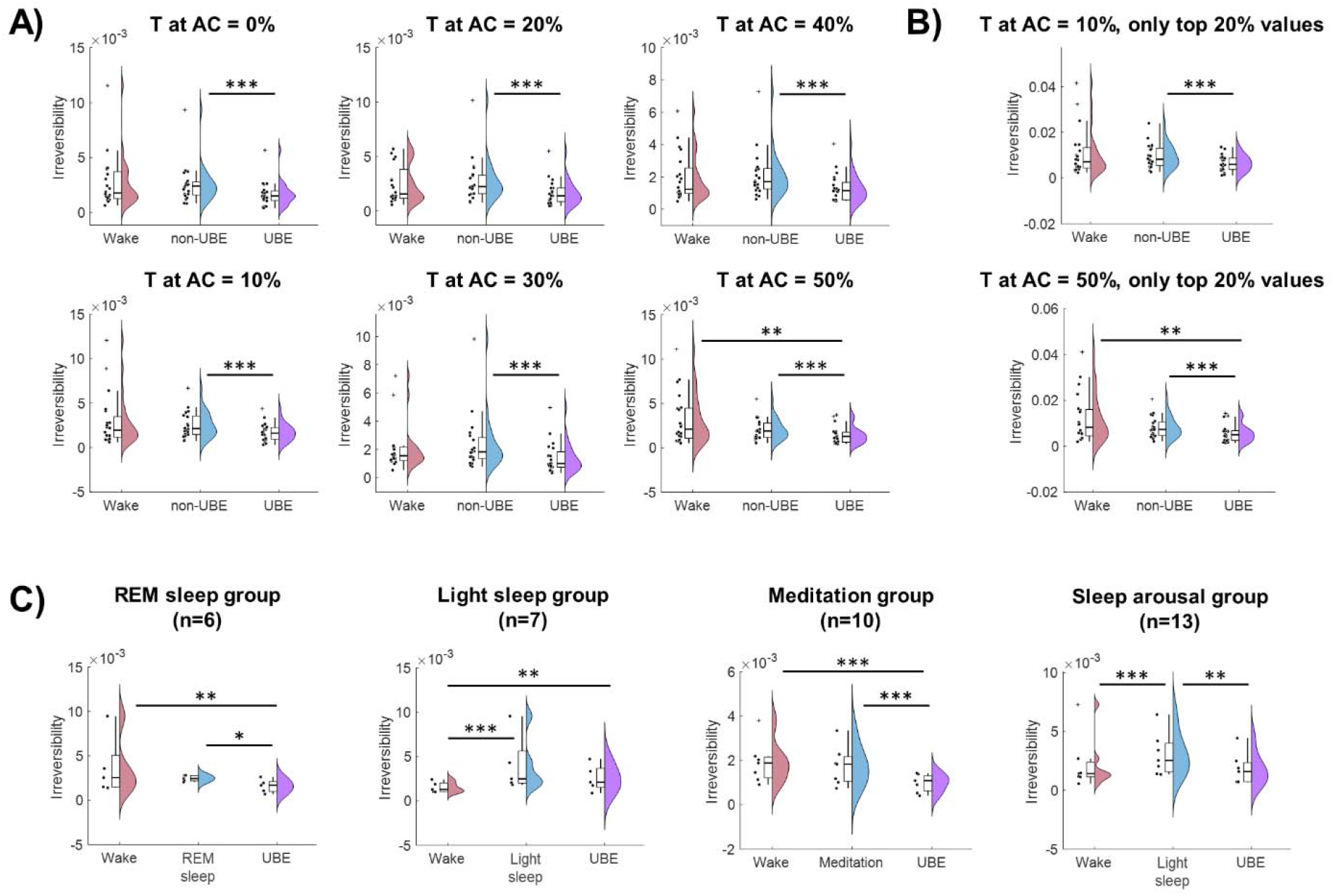
Sensitivity analyses of broadband EEG irreversibility. A) Time lag T was varied from 0% to 10%, 20%, 30%, 40% and 50% of the maximum autocorrelation (AC). B) Only the top 20% irreversibility values were considered when T was set at either AC = 10% or 50%. C) The dataset was split into groups according to the conscious stage during which UBEs were reported; SLEEP: ‘REM sleep’ group (n=6 UBEs, N=5 subjects) and ‘Light sleep’ group (n=7, N=5); WAKEFULNESS: ‘Meditation’ group (n=10, N=7) and ‘Sleep arousal’ group (n=13, N=7). Dots indicate subject-level means for irreversibility, while statistical comparisons with generalized linear mixed models were performed on individual trials following a within-subject design. Significance of model comparisons: *** p < 0.001, ** p < 0.01, * p < 0.05.

### Frequency-band EEG irreversibility

Next, we examined the contribution of multiple frequency bands to the overall brain irreversibility and hierarchy (**Figure S1**). For this purpose, the spectrum was divided into six conventional bands: delta (1-4 Hz), theta (4-8 Hz), alpha (8-12 Hz), sigma (12-15 Hz), beta (15-30 Hz), and gamma (30-45 Hz). Irreversibility decreased in UBEs compared to non-UBEs only in the alpha band (FDR-adjusted p < 0.05) and compared to wakefulness in the sigma band (FDR-adjusted p < 0.05). However, no significant differences were observed in hierarchy measures, nor when restricting the analysis to UBEs involving out-of-body experiences and feelings of elevation.

Additional sensitivity analyses (**Figure S2**) indicated that effects in specific frequency bands were largely consistent when considering only the top 30% irreversibility values, but strongly depended on the choice of T. For example, when T was set at AC = 50%, irreversibility significantly decreased in the alpha band in UBEs compared to wakefulness (FDR-adjusted p < 0.05) and significantly increased in the gamma band in UBEs compared to non-UBEs (FDR-adjusted p < 0.001). However, the differences originally observed when T was set at AC = 10% were no longer present in the alpha and sigma bands (see Supplementary Tables for detailed statistics).

### EEG complexity and spectral power

To complement the irreversibility analyses, we examined the same UBE dataset using two complementary approaches: (1) LZ complexity, which quantifies EEG signal diversity; (2) Multitaper spectral power analysis, which characterizes the distribution of oscillatory EEG activity across frequency bands (results previously reported in Campillo-Ferrer *et al*., 2026).

Complexity analyses (**Figure 5A**) revealed that non-UBE episodes show significantly lower global LZ complexity than both UBEs and wakefulness (FDR-adjusted p < 0.001). When restricting the analysis to UBEs involving out-of-body experiences and feelings of elevation, global LZ complexity remained significantly lower in non-UBEs compared to UBEs (FDR-adjusted p < 0.001), while no differences were observed between non-UBEs and wakefulness (FDR-adjusted p = 0.27). Likewise, no significant differences were found between wakefulness and UBEs in either the full (FDR-adjusted p = 0.93) or the restricted out-of-body sample (FDR-adjusted p = 0.54). In contrast, spectral power analyses (**Figure 5B**) characterized wakefulness as an activated EEG state, with increased power in higher frequencies (alpha, beta, and gamma) and decreased power in lower frequencies (delta and theta) compared to non-UBEs. UBEs showed an intermediate profile between wakefulness and non-UBEs, with decreased alpha and increased delta and theta activity relative to wakefulness, as well as reduced delta and theta and increased beta and gamma activity relative to non-UBEs (for detailed statistics see Campillo-Ferrer et al., 2026).

**Figure 5.**
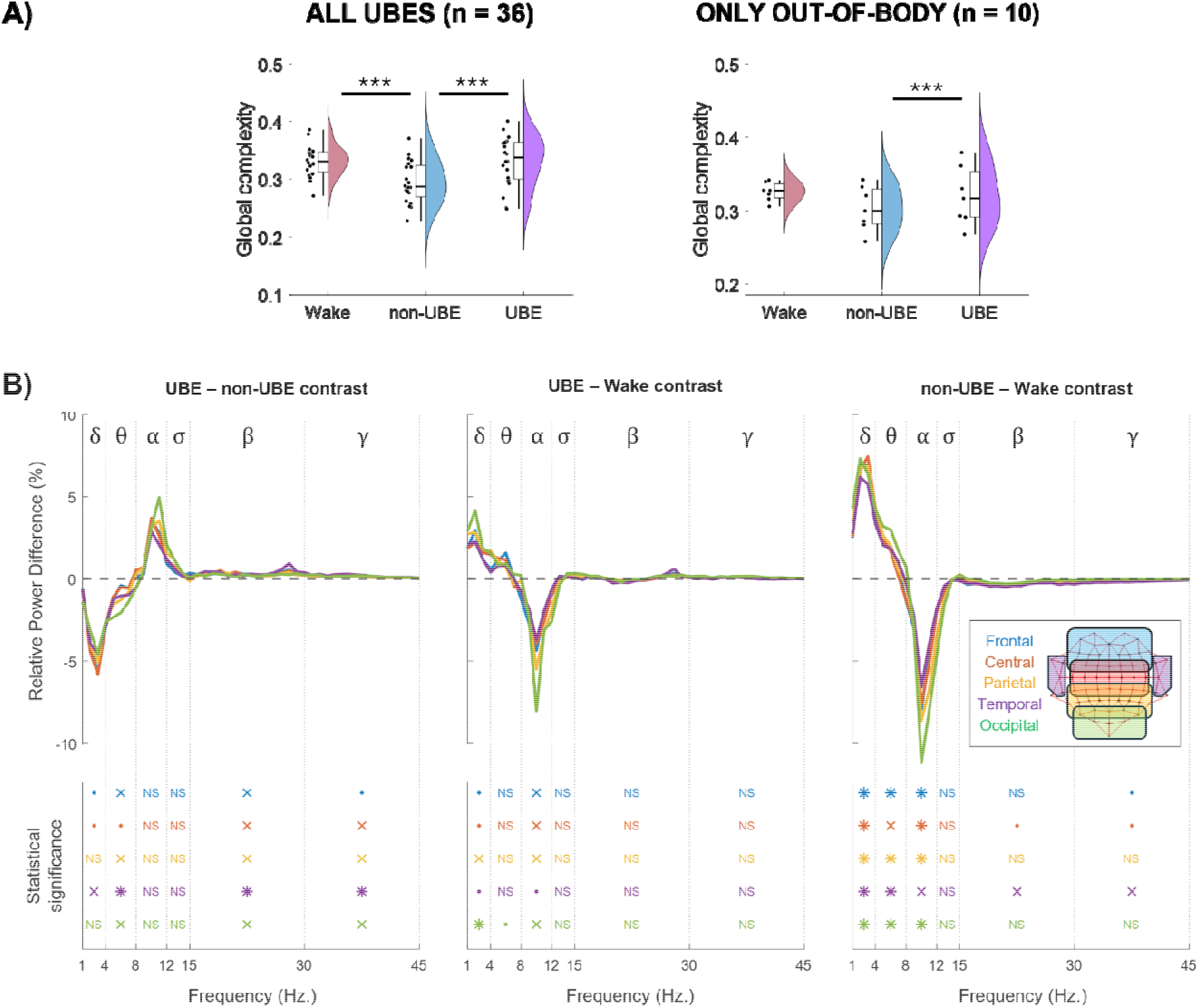
Complementary EEG analyses. A) Global LZ complexity across conditions in the full UBE sample (left, n=36 UBEs, N=20 subjects) and in UBEs involving out-of-body experiences and/or elevated self-location experiences (right, n=10 UBEs, N=7 subjects). Dots indicate subject-level means for visualization. Statistical comparisons with generalized linear mixed models were performed on individual trials and following a within-subject design. Significance of model comparisons: *** p < 0.001, ** p < 0.01, * p < 0.05. B) Relative power difference (%) for each region of interest: frontal (blue), central (red), parietal (yellow), temporal (purple), occipital (green), across frequency bands: delta (1-4 Hz), theta (4-8 Hz), alpha (8-12 Hz), sigma (12-15 Hz), beta (15-30 Hz), and gamma (30-45 Hz), and for each contrast: UBE vs. non-UBE (left), UBE vs. Wake (centre), non-UBE vs. Wake (right). The mean relative power was subtracted between conditions for visualization (e.g., UBE mean - non-UBE mean). Statistical comparisons with generalized linear mixed models were performed on individual trials using the original (non-subtracted) data and following a within-subject design. Significance of model comparisons: * p < 0.001, × p < 0.01, ▪ p < 0.05, NS = not significant.

**Table 1** summarizes the main results obtained from irreversibility, hierarchy, complexity, and spectral analyses. As illustrated, EEG activation derived from spectral analyses followed a graded pattern: high during wakefulness, low during non-UBEs, and intermediate during UBEs, characterizing UBEs as a hybrid state between wakefulness and non-UBEs. In contrast, irreversibility and hierarchy were high during wakefulness and non-UBEs but significantly reduced during UBEs, indicating a decrease of temporal directionality or asymmetry in this experimental condition, together with a decrease of cortical hierarchical organization. Complexity, on the other hand, remained high in both wakefulness and UBEs but low during non-UBEs, reflecting preserved signal diversity during UBEs.

**Table 1:**
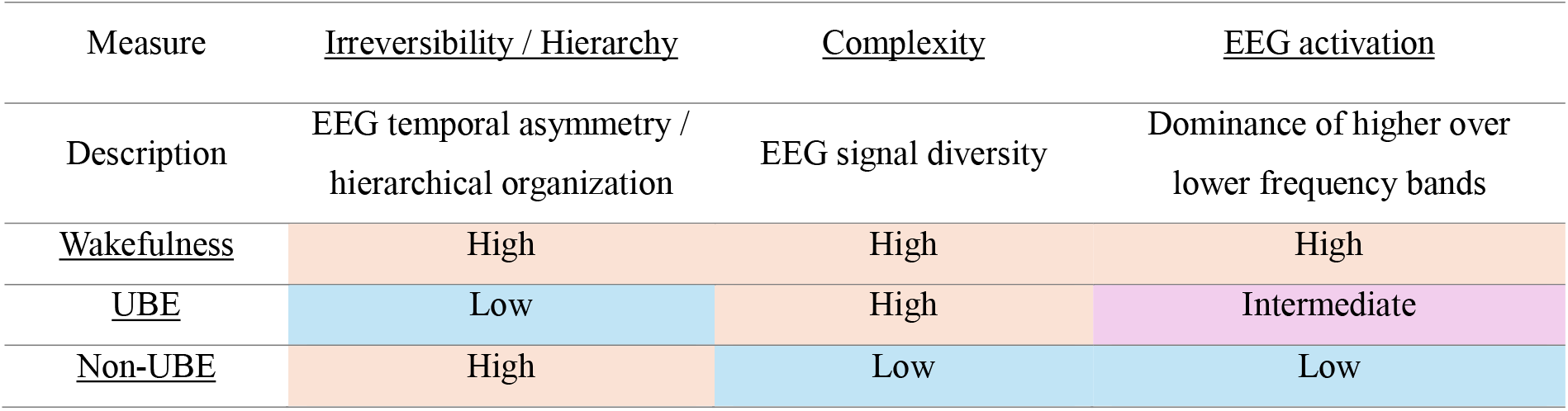
EEG signatures of wakefulness, UBEs and non-UBEs.

## Discussion

Our findings show that UBEs are associated with a transient reduction in EEG irreversibility and hierarchical differentiation compared to non-UBEs and wakefulness, a pattern observed consistently across multiple states of consciousness. In contrast, global LZ complexity remains high during UBEs and at levels comparable to wakefulness. These results are robust across sensitivity analyses and generalized linear mixed models (GLMMs) at the single-trial level. Importantly, the use of GLMMs allowed us to study UBEs across different states of consciousness within a unified framework, including experiences reported during REM sleep, light sleep, meditation, and sleep arousals. Supplementary frequency-band analyses further suggest that these patterns primarily manifest at a broadband and global scale.

### UBEs are associated with a global flattening of brain hierarchy

Conceptually, irreversibility and hierarchy capture complementary dimensions of temporal asymmetry in neural activity. Irreversibility measures the overall magnitude of temporal asymmetry in the neural dynamics (i.e., whether information carried by brain activity flows more in one direction than the reverse), whereas hierarchy quantifies how unevenly that asymmetry is distributed across cortical regions (i.e., whether all regions contribute similarly to temporal asymmetry or not). Reductions in irreversibility and hierarchy observed during UBEs may thus reflect a temporary flattening of cortical hierarchy relative to non-UBEs (**Figure 3**). These results are consistent with previous findings showing that irreversibility is typically highest during wakefulness and decreases in states of reduced consciousness like deep sleep or coma (Berjaga-Buisan *et al*., 2025; Camassa *et al*., 2024; Deco *et al*., 2022; G-Guzmán *et al*., 2023; Idesis *et al*., 2024; Sanz Perl *et al*., 2021). Reduced irreversibility has also been observed during movie watching compared to both rest and task engagement (Kringelbach *et al*., 2023), suggesting that UBEs may not necessarily reflect a deeper unconscious state, but rather a more passive mode of brain activity with diminished computational demands.

Reductions in irreversibility were also observed in UBEs compared to resting wakefulness, but these effects were dependent on specific parameter choices (**Figures 4A** and **4B**). The pattern became clearer when UBEs were analyzed separately according to their state of consciousness (**Figure 4C**), revealing significant differences in all groups except for the ‘sleep arousal’ group. Importantly, our findings suggest that these effects reflect changes specifically associated with UBEs, rather than intrinsic properties of their respective baseline states. In this context, non-UBE conditions offer a useful basis for comparison, as they either resembled wakefulness or showed opposite effects to UBEs. Specifically, no significant differences in irreversibility were found when comparing wakefulness with non-UBEs during meditation or REM sleep (**Figure 4C**, ‘REM sleep’ and ‘Meditation’ groups). Likewise, although non-UBEs during light sleep differed significantly from wakefulness, the direction of this effect was opposite to that observed during UBEs in the same state (**Figure 4C**, ‘Light sleep’ and ‘Sleep arousal’ groups). In this context, it is worth noting that comparisons between wakefulness and non-UBE conditions are consistent with both prior literature and the specific context of our dataset. First, minimal differences observed between wakefulness and meditation in the non-UBE condition likely reflect the limited meditation experience of our participants, while the use of eyes-closed wakefulness under mild sleep deprivation (2–3 hours) may have attenuated contrasts with sleep. Second, previous work in the ferret cortex has shown that irreversibility during REM sleep can approach levels observed in wakefulness (Idesis et al., 2024), although direct comparisons between wakefulness and light sleep remain unexplored.

Taken together, these findings suggest that UBEs are associated with a robust decrease in the directedness of neural dynamics and hierarchical differentiation compared to both non-UBEs and wakefulness, while non-UBE conditions largely resembled wakefulness or showed effects in the opposite direction. Further analyses suggest that these effects emerge only at a broadband and global scale. For example, analyses focused on canonical frequency bands (delta, theta, alpha, sigma, beta, and gamma) revealed no significant differences in hierarchy across conditions (**Figure S1**), while changes in irreversibility were highly sensitive to the choice of T (**Figure S2**). Thus, the reduction in irreversibility and hierarchy observed during UBEs cannot reliably be attributed to any specific frequency band, but may instead reflect a broadband (and potentially global) reorganization of brain dynamics. Consistent with this interpretation, topographic maps show that the same cortical regions contribute to irreversibility across all conditions, with UBEs differing mainly in the magnitude of irreversibility rather than its spatial distribution (**Figure 3**, bottom). However, future studies employing source reconstruction are needed to confirm these findings at the cortical level.

### UBEs are associated with increased global complexity, resembling the psychedelic state

Analyses of complexity offer a complementary perspective on the brain dynamics associated with UBEs. Whereas irreversibility and hierarchy decrease during these states, global LZ complexity shows the opposite pattern: it increases compared to non-UBEs, and remains at levels similar to wakefulness (**Figure 5**, top). Decreases in irreversibility have typically been associated with reduced complexity and lower levels of consciousness in the literature. For instance, reductions in both irreversibility and LZ complexity have been observed in coma, deep sleep, and disorders of consciousness (Aamodt *et al*., 2022; Camassa *et al*., 2024; Deco *et al*., 2022; G-Guzmán *et al*., 2023; Idesis *et al*., 2024; Liu *et al*., 2023; Schartner *et al*., 2015; Schartner, Pigorini, *et al*., 2017; Zhang *et al*., 2001). However, recent evidence suggests that these two metrics can dissociate: psychedelic states, long known for presenting elevated global complexity (Pallavicini *et al*., 2021; Schartner, Carhart-Harris, *et al*., 2017; Timmermann *et al*., 2019) can also exhibit decreased irreversibility compared to placebo (Barnett *et al*., 2020; Shinozuka *et al*., 2025). Additionally, one study measured both irreversibility and complexity in the same dataset, showing that high global complexity can coexist with low irreversibility in the psychedelic state (Vohryzek *et al*., 2025). The present study suggests that UBEs may exhibit a similar pattern, combining wake-like levels of global complexity with reduced irreversibility.

One possible interpretation for this combination is that UBEs reflect a state characterized by a rich variety of EEG patterns (high complexity), while organizing these patterns in a more flexible and less constrained manner (low irreversibility and hierarchy). Under these conditions, information may propagate through alternative pathways, potentially reflecting a state less constrained by top-down organization. A redistribution of information flow through atypical pathways could help explain the phenomenology of some UBEs analysed here, where participants perceived their body as a single “shape”, “contour”, or the laboratory room as an “immensely huge universe” (see phenomenological reports in Campillo-Ferrer *et al*., 2026). Interestingly, a similar interpretation has been proposed in studies of psychedelic-induced “ego-dissolution” states, where individuals may experience a sense of merging with their surroundings (Millière, 2017; Nour *et al*., 2016). These studies suggest that ego-dissolution is associated with a reorganization of brain connectivity, interpreted as a “collapse” of the brain’s normal hierarchical organization (Lebedev *et al*., 2015; Tagliazucchi *et al*., 2016).

A global reorganization of brain dynamics may similarly facilitate the occurrence of out-of-body experiences, which are also reported under psychedelics and other altered states (Herrero *et al*., 2025; Stoliker *et al*., 2025; Twemlow *et al*., 1982). In the present study, these experiences were analysed together with the full set of UBEs, but we also performed a separate analysis focusing exclusively on out-of-body experiences and feelings of elevation (n=10). Some of these experiences involved the sensation of the “soul” ascending and leaving the physical body, or the sensation of the body being “lifted” from a specific point in the chest. Overall, these experiences showed the same patterns observed in the full UBE sample, characterized by reduced broadband irreversibility and hierarchy compared to both non-UBEs and wakefulness (**Figure 3B**), increased global LZ complexity compared to non-UBEs (**Figure 5A**, right), and no significant frequency-band effects (**Figure S1B**). Additionally, the cortical regions contributing to irreversibility closely resembled those identified in the full UBE sample (**Figure 3B**, bottom). Taken together, these findings support the notion that out-of-body experiences may exist on a continuum, potentially sharing underlying brain dynamics with other phenomena such as ego-dissolution and vestibular-motor hallucinations (Blackmore, 1988; Campillo-Ferrer *et al*., 2024; Cheyne & Girard, 2009).

Crucially, these results can also be interpreted through a non-equilibrium thermodynamic perspective (Kringelbach *et al*., 2024). From this perspective, both UBEs and out-of-body experiences appear to emerge when the brain shifts slightly toward equilibrium, reflecting a state of consciousness in which information flow becomes less directional and more evenly distributed across brain regions. However, although proximity to equilibrium has traditionally been linked to diminished consciousness, participants in our study reported conscious awareness during UBEs in most cases (71%, see the “lucid insight” category in Campillo-Ferrer *et al*., 2026). Similarly, previous studies have reported reduced irreversibility during psychedelic states relative to placebo (Barnett *et al*., 2020; Shinozuka *et al*., 2025; Vohryzek *et al*., 2025), despite these states being associated with rich phenomenology and conscious awareness (Lawrence *et al*., 2022; Metastasio *et al*., 2025). These findings indicate that a shift toward equilibrium does not necessarily imply reduced consciousness, but may rather reflect a reconfiguration of brain dynamics that supports conscious experience in atypical ways (Huang *et al*., 2023).

### UBEs beyond intermediate states of consciousness

Finally, it is important to emphasize that measures of irreversibility, hierarchy, and complexity provide unique insights into the EEG correlates of UBEs that go beyond those offered by conventional spectral analyses. While multitaper spectral approaches characterized UBEs as intermediate states between wakefulness and sleep (see **Figure 5B**), analyses of irreversibility, hierarchy, and complexity instead suggest that UBEs constitute a distinct state of consciousness different from both. Preliminary evidence from principal component analyses (PCA) of EEG further supports this view, showing that out-of-body experiences and sleep paralysis episodes form non-overlapping clusters in the PCA space (Herrero *et al*., 2025), instead of simply occupying an intermediate position between canonical states like wakefulness and sleep, as suggested by other studies (Campillo-Ferrer *et al*., 2026; Mainieri *et al*., 2021; Terzaghi *et al*., 2012).

Overall, these results suggest that UBEs constitute a distinct state of consciousness, highlighting novel patterns of neural organization and providing a unique window into the dynamics of conscious experience and bodily perception. Future studies using whole-brain modelling and causal perturbations may help clarify the causal mechanisms underlying UBEs, defining the boundaries between different brain states and exploring how the awake brain could be guided into (or out of) UBEs and out-of-body experiences (Dagnino *et al*., 2024; Deco *et al*., 2019). Such insights could have important clinical applications, for example, to enhance the understanding and management of schizophrenia, death anxiety, and other mental health conditions (Bünning & Blanke, 2005; Shaw *et al*., 2023).

### Limitations

This study has several limitations that largely mirror those of the original work (see Campillo-Ferrer *et al*., 2026). Although micro-phenomenological interviews provide detailed insights, they remain subjective and may not fully capture all UBEs or their precise timing. In particular, the selection of 5-second EEG segments to characterize each UBE report may introduce some temporal imprecision. Furthermore, dream reports were not collected for non-UBE periods and the induction protocol included sensory stimulation, so these episodes may have contained unreported UBEs, dreams that were not consciously remembered, or brief sleep disruptions potentially affecting the data.

## Conclusion

Our findings suggest that brain activity during UBEs is marked by a transient flattening of global hierarchical dynamics relative to both non-UBEs and wakefulness, while preserving high informational richness equivalent to wakefulness. This combination resembles the neural dynamics observed in psychedelic states and may potentially reflect a state of consciousness unconstrained by higher-level mental constructs, where bottom-up information propagates more flexibly through alternative neural pathways.

## Author contributions

Conceptualization and methodology: TC-F, TB-B, and GD. Sleep scoring: TC-F, AI, RC, MV, and LN. EEG preprocessing: TC-F and RO. EEG analysis, statistical testing, and visualization: TC-F and TB-B. Writing – original draft: TC-F and TB-B. Writing – review and editing: all authors. Project supervision: GD. Project administration: TC-F and GD.

## Funding

This project was funded by the BIAL Foundation grants programme for scientific research 2024 - 81/24.

## Data availability

The data that support the findings of this study are openly available in the Radboud University repository at https://doi.org/10.34973/50bx-st45.

## Code availability

The open-source MATLAB code used to apply the INSIDEOUT method can be found at https://github.com/tecferrer/UBE-INSIDEOUT-code.

## Declaration of competing interest

The authors declare that they have no known competing financial interests or personal relationships that could have appeared to influence the work reported in this paper.

## Notes

### Competing Interest Statement

The authors have declared no competing interest.

## References

Aamodt, A., Sevenius Nilsen, A., Markhus, R., Kusztor, A., HasanzadehMoghadam, F., Kauppi, N., Thürer, B., Storm, J. F., & Juel, B. E. (2022). EEG Lempel-Ziv complexity varies with sleep stage, but does not seem to track dream experience. Frontiers in Human Neuroscience, 16, 987714. 10.3389/fnhum.2022.987714

Alcaraz-Sánchez, A., Demšar, E., Campillo-Ferrer, T., & Torres-Platas, S. G. (2022). Nothingness Is All There Is: An Exploration of Objectless Awareness During Sleep. Frontiers in Psychology, 13, 901031. 10.3389/fpsyg.2022.901031

Barnett, L., Muthukumaraswamy, S. D., Carhart-Harris, R. L., & Seth, A. K. (2020). Decreased directed functional connectivity in the psychedelic state. NeuroImage, 209, 116462. 10.1016/j.neuroimage.2019.116462

Berjaga-Buisan, T., Monti, J. M., Cortada, M., Colombo, M. A., Geli, S. M., Gaglioti, G., Sarasso, S., Kringelbach, M. L., Corbetta, M., Sanchez-Vives, M. V., Massimini, M., Perl, Y. S., & Deco, G. (2025). Thermodynamics of consciousness: A non-invasive perturbational framework. bioRxiv, 2025.12.09.691422. 10.64898/2025.12.09.691422

Berkovich-Ohana, A., Dor-Ziderman, Y., Glicksohn, J., & Goldstein, A. (2013). Alterations in the sense of time, space, and body in the mindfulness-trained brain: A neurophenomenologically-guided MEG study. Frontiers in Psychology, 4, 912. 10.3389/fpsyg.2013.00912

Berry, R. B., Quan, S. F., Abreu, A. R., Bibbs, M. L., DelRosso, L., Harding, S. M., Mao, M.-M., Plante, D. T., Pressman, M. R., Troester, M. M., & Vaughnm, B. V. (2020). The AASM Manual for the Scoring of Sleep and Associated Events: Rules, Terminology and Technical Specifications. *Version 2.6.* American Academy of Sleep Medicine.

Blackmore, S. J. (1988). A Theory of Lucid Dreams and OBEs. In J. Gackenbach & S. LaBerge (Eds.), Conscious Mind, Sleeping Brain: Perspectives on Lucid Dreaming (pp. 373–387). Springer New York. 10.1007/978-1-4757-0423-5_16

Blanke, O., & Arzy, S. (2005). The out-of-body experience: Disturbed self-processing at the temporo-parietal junction. *The Neuroscientist: A Review Journal Bringing Neurobiology*, Neurology and Psychiatry, 11(1), 16–24. 10.1177/1073858404270885

Blanke, O., Landis, T., Spinelli, L., & Seeck, M. (2004). Out-of-body experience and autoscopy of neurological origin. Brain: A Journal of Neurology, 127(Pt 2), 243–258. 10.1093/brain/awh040

Blanke, O., Ortigue, S., Landis, T., & Seeck, M. (2002). Stimulating illusory own-body perceptions. Nature, 419(6904), 269–270. 10.1038/419269a

Blanke, O., Perrig, S., Thut, G., Landis, T., & Seeck, M. (2000). Simple and complex vestibular responses induced by electrical cortical stimulation of the parietal cortex in humans. Journal of Neurology, Neurosurgery, and Psychiatry, 69(4), 553–556. 10.1136/jnnp.69.4.553

Bünning, S., & Blanke, O. (2005). The out-of body experience: Precipitating factors and neural correlates. Progress in Brain Research, 150, 331–350. 10.1016/s0079-6123(05)50024-4

Camassa, A., Torao-Angosto, M., Manasanch, A., Kringelbach, M. L., Deco, G., & Sanchez-Vives, M. V. (2024). The temporal asymmetry of cortical dynamics as a signature of brain states. Scientific Reports, 14(1), 24271. 10.1038/s41598-024-74649-1

Campillo-Ferrer, T., Alcaraz-Sánchez, A., Demšar, E., Wu, H.-P., Dresler, M., Windt, J., & Blanke, O. (2024). Out-of-body experiences in relation to lucid dreaming and sleep paralysis: A theoretical review and conceptual model. Neuroscience and Biobehavioral Reviews, 163, 105770. 10.1016/j.neubiorev.2024.105770

Campillo-Ferrer, T., Alcaraz-Sánchez, A., & Torres-Platas, S. G. (2025). Exploring “lucid sleep” and altered states of consciousness using meditation and visual stimulation: A case series study. Philosophy and the Mind Sciences, 5. 10.33735/phimisci.2024.10227

Campillo-Ferrer, T., Iadarola, A., Cordani, R., Veneruso, M., Demirel, C., Nobili, L., & Oostenveld, R. (2026). Facilitating unusual bodily experiences and out-of-body experiences across wakefulness and sleep: A high-density EEG and neurophenomenology study. Consciousness and Cognition, 139, 104002. 10.1016/j.concog.2026.104002

Cheyne, J. A. (2003). Sleep paralysis and the structure of waking-nightmare hallucinations. Dreaming, 13(3), 163–179. 10.1023/A:1025373412722

Cheyne, J. A., & Girard, T. A. (2009). The body unbound: Vestibular-motor hallucinations and out-of-body experiences. Cortex; a Journal Devoted to the Study of the Nervous System and Behavior, 45(2), 201–215. 10.1016/j.cortex.2007.05.002

Cheyne, J. A., Rueffer, S. D., & Newby-Clark, I. R. (1999). Hypnagogic and hypnopompic hallucinations during sleep paralysis: Neurological and cultural construction of the night-mare. Consciousness and Cognition, 8(3), 319–337. 10.1006/ccog.1999.0404

Cruzat, J., Herzog, R., Prado, P., Sanz-Perl, Y., Gonzalez-Gomez, R., Moguilner, S., Kringelbach, M. L., Deco, G., Tagliazucchi, E., & Ibañez, A. (2023). Temporal Irreversibility of Large-Scale Brain Dynamics in Alzheimer’s Disease. The Journal of Neuroscience: The Official Journal of the Society for Neuroscience, 43(9), 1643–1656. 10.1523/jneurosci.1312-22.2022

Dagnino, P. C., Escrichs, A., López-González, A., Gosseries, O., Annen, J., Sanz Perl, Y., Kringelbach, M. L., Laureys, S., & Deco, G. (2024). Re-awakening the brain: Forcing transitions in disorders of consciousness by external in silico perturbation. PLoS Computational Biology, 20(5), e1011350. 10.1371/journal.pcbi.1011350

De Ridder, D., Van Laere, K., Dupont, P., Menovsky, T., & Van de Heyning, P. (2007). Visualizing out-of-body experience in the brain. The New England Journal of Medicine, 357(18), 1829–1833. 10.1056/NEJMoa070010

Deco, G., Cruzat, J., Cabral, J., Tagliazucchi, E., Laufs, H., Logothetis, N. K., & Kringelbach, M. L. (2019). Awakening: Predicting external stimulation to force transitions between different brain states. Proceedings of the National Academy of Sciences of the United States of America, 116(36), 18088–18097. 10.1073/pnas.1905534116

Deco, G., Sanz Perl, Y., Bocaccio, H., Tagliazucchi, E., & Kringelbach, M. L. (2022). The INSIDEOUT framework provides precise signatures of the balance of intrinsic and extrinsic dynamics in brain states. Communications Biology, 5(1), 572. 10.1038/s42003-022-03505-7

Deco, G., Sanz Perl, Y., de la Fuente, L., Sitt, J. D., Yeo, B. T. T., Tagliazucchi, E., & Kringelbach, M. L. (2023). The arrow of time of brain signals in cognition: Potential intriguing role of parts of the default mode network. Network Neuroscience, 7(3), 966–998. 10.1162/netn_a_00300

Demirel, Ç., Gott, J., Appel, K., Lüth, K., Fischer, C., Raffaelli, C., Westner, B. U., Wang, X., Zavecz, Z., Steiger, A., Erlacher, D., LaBerge, S., Mota-Rolim, S. A., Ribeiro, S., Zeising, M., Adelhöfer, N., & Dresler, M. (2025). Electrophysiological Correlates of Lucid Dreaming: Sensor and Source Level Signatures. The Journal of Neuroscience: The Official Journal of the Society for Neuroscience, 45(20), e2237242025. 10.1523/JNEUROSCI.2237-24.2025

Dor-Ziderman, Y., Ataria, Y., Fulder, S., Goldstein, A., & Berkovich-Ohana, A. (2016). Self-specific processing in the meditating brain: A MEG neurophenomenology study. Neuroscience of Consciousness, 2016(1), niw019. 10.1093/nc/niw019

Eddington, A. S. (1928). The Nature of the Physical World. New York : The Macmillan Company.

Faul, S. (2025, October). Kolmogorov Complexity. MATLAB Central File Exchange. https://www.mathworks.com/matlabcentral/fileexchange/6886-kolmogorov-complexity

Feng, E. H., & Crooks, G. E. (2008). Length of Time’s Arrow. Physical Review Letters, 101(9), 090602. 10.1103/PhysRevLett.101.090602

G-Guzmán, E., Perl, Y. S., Vohryzek, J., Escrichs, A., Manasova, D., Türker, B., Tagliazucchi, E., Kringelbach, M., Sitt, J. D., & Deco, G. (2023). The lack of temporal brain dynamics asymmetry as a signature of impaired consciousness states. Interface Focus, 13(3), 20220086. 10.1098/rsfs.2022.0086

Herrero, N. L., Corfdir, Y., Vázquez-Chenlo, A. A., Capurro, L., & Forcato, C. (2025). Exploratory study of non-ordinary states of consciousness during sleep show distinct electrophysiological features from wakefulness and canonical sleep stages. Scientific Reports, 15(1), 33586. 10.1038/s41598-025-18748-7

Huang, Z., Mashour, G. A., & Hudetz, A. G. (2023). Functional geometry of the cortex encodes dimensions of consciousness. Nature Communications, 14(1), 72. 10.1038/s41467-022-35764-7

Idesis, S., Geli, S., Faskowitz, J., Vohryzek, J., Sanz Perl, Y., Pieper, F., Galindo-Leon, E., Engel, A. K., & Deco, G. (2024). Functional hierarchies in brain dynamics characterized by signal reversibility in ferret cortex. PLoS Computational Biology, 20(1), e1011818. 10.1371/journal.pcbi.1011818

Ionta, S., Heydrich, L., Lenggenhager, B., Mouthon, M., Fornari, E., Chapuis, D., Gassert, R., & Blanke, O. (2011). Multisensory mechanisms in temporo-parietal cortex support self-location and first-person perspective. Neuron, 70(2), 363–374. 10.1016/j.neuron.2011.03.009

Karvelis, P. (2025, July). Daviolinplot—Violin and raincloud plots. GitHub. https://github.com/povilaskarvelis/DataViz/releases/tag/v3.2.7

Kaspar, F., & Schuster, H. G. (1987). Easily calculable measure for the complexity of spatiotemporal patterns. *Physical Review. A*, General Physics, 36(2), 842–848. 10.1103/physreva.36.842

Kringelbach, M. L., Perl, Y. S., Tagliazucchi, E., & Deco, G. (2023). Toward naturalistic neuroscience: Mechanisms underlying the flattening of brain hierarchy in movie-watching compared to rest and task. Science Advances, 9(2), eade6049. 10.1126/sciadv.ade6049

Kringelbach, M. L., Sanz Perl, Y., & Deco, G. (2024). The Thermodynamics of Mind. Trends in Cognitive Sciences, 28(6), 568–581. 10.1016/j.tics.2024.03.009

LaBerge, S., Nagel, L., Dement, W., & Zarcone, V. (1981). Lucid dreaming verified by volitional communication during REM sleep. Perceptual and Motor Skills, 52(3), 727–732. 10.2466/pms.1981.52.3.727

Lawrence, D. W., Carhart-Harris, R., Griffiths, R., & Timmermann, C. (2022). Phenomenology and content of the inhaled N, N-dimethyltryptamine (N, N-DMT) experience. Scientific Reports, 12(1), 8562. 10.1038/s41598-022-11999-8

Lebedev, A. V., Lövdén, M., Rosenthal, G., Feilding, A., Nutt, D. J., & Carhart-Harris, R. L. (2015). Finding the self by losing the self: Neural correlates of ego-dissolution under psilocybin. Human Brain Mapping, 36(8), 3137–3153. 10.1002/hbm.22833

Lempel, A., & Ziv, J. (1976). On the complexity of finite sequence. IEEE Transactions on Information Theory, 22(1), 75–81. 10.1109/TIT.1976.1055501

Lenggenhager, B., Halje, P., & Blanke, O. (2011). Alpha band oscillations correlate with illusory self-location induced by virtual reality. The European Journal of Neuroscience, 33(10), 1935–1943. 10.1111/j.1460-9568.2011.07647.x

Levitan, L., LaBerge, S., DeGracia, D. J., & Zimbardo, P. G. (1999). Out-of-body experiences, dreams, and REM sleep. Sleep and Hypnosis, 1(3), 186–196.

Liu, Y., Zeng, W., Pan, N., Xia, X., Huang, Y., & He, J. (2023). EEG complexity correlates with residual consciousness level of disorders of consciousness. BMC Neurology, 23(1), 140. 10.1186/s12883-023-03167-w

Mainieri, G., Maranci, J. B., Champetier, P., Leu-Semenescu, S., Gales, A., Dodet, P., & Arnulf, I. (2021). Are sleep paralysis and false awakenings different from REM sleep and from lucid REM sleep? A spectral EEG analysis. Journal of Clinical Sleep Medicine: JCSM: Official Publication of the American Academy of Sleep Medicine, 17(4), 719–727. 10.5664/jcsm.9056

Martial, C., Cassol, H., Slater, M., Bourdin, P., Mensen, A., Oliva, R., Laureys, S., & Núñez, P. (2023). Electroencephalographic Signature of Out-of-Body Experiences Induced by Virtual Reality: A Novel Methodological Approach. Journal of Cognitive Neuroscience, 35(9), 1410–1422. 10.1162/jocn_a_02011

Marzano, C., Moroni, F., Gorgoni, M., Nobili, L., Ferrara, M., & De Gennaro, L. (2013). How we fall asleep: Regional and temporal differences in electroencephalographic synchronization at sleep onset. Sleep Medicine, 14(11), 1112–1122. 10.1016/j.sleep.2013.05.021

McCreery, C., & Claridge, G. (1996). A study of hallucination in normal subjects—II. Electrophysiological data. Personality and Individual Differences, 21(5), 749–758. 10.1016/0191-8869(96)00116-X

Metastasio, A., Prevete, E., Venturini, S., Garofalo, A., Cecconello, B., De Pisapia, N., & Corazza, O. (2025). The phenomenology of psilocybin: Transformative insights for research and clinical practice. Frontiers in Psychology, 16, 1455902. 10.3389/fpsyg.2025.1455902

Millière, R. (2017). Looking for the Self: Phenomenology, Neurophysiology and Philosophical Significance of Drug-induced Ego Dissolution. Frontiers in Human Neuroscience, 11, 245. 10.3389/fnhum.2017.00245

Nour, M. M., Evans, L., Nutt, D., & Carhart-Harris, R. L. (2016). Ego-Dissolution and Psychedelics: Validation of the Ego-Dissolution Inventory (EDI). Frontiers in Human Neuroscience, 10, 269. 10.3389/fnhum.2016.00269

Oostenveld, R., Fries, P., Maris, E., & Schoffelen, J. M. (2011). FieldTrip: Open source software for advanced analysis of MEG, EEG, and invasive electrophysiological data. Computational Intelligence and Neuroscience, 2011, 156869. 10.1155/2011/156869

Pallavicini, C., Cavanna, F., Zamberlan, F., Fuente, L. A. de la, Ilksoy, Y., Perl, Y. S., Arias, M., Romero, C., Carhart-Harris, R., Timmermann, C., & Tagliazucchi, E. (2021). Neural and subjective effects of inhaled N,N-dimethyltryptamine in natural settings. Journal of Psychopharmacology, 35(4), 406–420. 10.1177/0269881120981384

Petitmengin, C. (2006). Describing one’s subjective experience in the second person: An interview method for the science of consciousness. Phenomenology and the Cognitive Sciences, 5(3–4), 229–269. 10.1007/s11097-006-9022-2

Petitmengin, C., Remillieux, A., & Valenzuela-Moguillansky, C. (2019). Discovering the structures of lived experience: Towards a micro-phenomenological analysis method. Phenomenology and the Cognitive Sciences, 18(4), 691–730. 10.1007/s11097-018-9597-4

Rabeyron, T., & Caussie, S. (2016). Clinical aspects of Out-of-Body Experiences: Trauma, reflexivity and symbolisation. *L’**É*volution Psychiatrique, 81(4), e53–e71. 10.1016/j.evopsy.2016.09.002

Sanz Perl, Y., Bocaccio, H., Pallavicini, C., Pérez-Ipiña, I., Laureys, S., Laufs, H., Kringelbach, M., Deco, G., & Tagliazucchi, E. (2021). Nonequilibrium brain dynamics as a signature of consciousness. Physical Review. E, 104(1–1), 014411. 10.1103/PhysRevE.104.014411

Schartner, M. M., Carhart-Harris, R. L., Barrett, A. B., Seth, A. K., & Muthukumaraswamy, S. D. (2017). Increased spontaneous MEG signal diversity for psychoactive doses of ketamine, LSD and psilocybin. Scientific Reports, 7(1), 46421. 10.1038/srep46421

Schartner, M. M., Pigorini, A., Gibbs, S. A., Arnulfo, G., Sarasso, S., Barnett, L., Nobili, L., Massimini, M., Seth, A. K., & Barrett, A. B. (2017). Global and local complexity of intracranial EEG decreases during NREM sleep. Neuroscience of Consciousness, 2017(1), niw022. 10.1093/nc/niw022

Schartner, M. M., Seth, A., Noirhomme, Q., Boly, M., Bruno, M.-A., Laureys, S., & Barrett, A. (2015). Complexity of Multi-Dimensional Spontaneous EEG Decreases during Propofol Induced General Anaesthesia. PloS One, 10(8), e0133532. 10.1371/journal.pone.0133532

Seif, A., Hafezi, M., & Jarzynski, C. (2021). Machine learning the thermodynamic arrow of time. Nature Physics, 17(1), 105–113. 10.1038/s41567-020-1018-2

Shaw, J., Gandy, S., & Stumbrys, T. (2023). Transformative effects of spontaneous out of body experiences in healthy individuals: An interpretative phenomenological analysis. Psychology of Consciousness: Theory, Research, and Practice, Advance online publication. 10.1037/cns0000324

Shinozuka, K., Tewarie, P. K. B., Luppi, A., Lynn, C., Roseman, L., Muthukumaraswamy, S., Nutt, D. J., Carhart-Harris, R., Deco, G., & Kringelbach, M. L. (2025). LSD flattens the hierarchy of directed information flow in fast whole-brain dynamics. Imaging Neuroscience, 3, imag_a_00420. 10.1162/imag_a_00420

Siclari, F., Bernardi, G., Riedner, B. A., LaRocque, J. J., Benca, R. M., & Tononi, G. (2014). Two distinct synchronization processes in the transition to sleep: A high-density electroencephalographic study. Sleep, 37(10), 1621–1637. 10.5665/sleep.4070

Smith, A. M., & Messier, C. (2014). Voluntary Out-of-Body Experience: An fMRI Study. Frontiers in Human Neuroscience, 8, 70. 10.3389/fnhum.2014.00070

Stoliker, D., Bernasconi, F., Blanke, O., & Razi, A. (2025). Psilocybin Modulates TPJ Effective Connectivity during Out-of-Body Experiences. MedRxiv Preprint, 2025.06.24.25330245. 10.1101/2025.06.24.25330245

Tagliazucchi, E., Roseman, L., Kaelen, M., Orban, C., Muthukumaraswamy, S. D., Murphy, K., Laufs, H., Leech, R., McGonigle, J., Crossley, N., Bullmore, E., Williams, T., Bolstridge, M., Feilding, A., Nutt, D. J., & Carhart-Harris, R. (2016). Increased Global Functional Connectivity Correlates with LSD-Induced Ego Dissolution. Current Biology: CB, 26(8), 1043–1050. 10.1016/j.cub.2016.02.010

Takeuchi, T., Fukuda, K., Sasaki, Y., Inugami, M., & Murphy, T. I. (2002). Factors related to the occurrence of isolated sleep paralysis elicited during a multi-phasic sleep-wake schedule. Sleep, 25(1), 89–96. 10.1093/sleep/25.1.89

Takeuchi, T., Miyasita, A., Sasaki, Y., Inugami, M., & Fukuda, K. (1992). Isolated sleep paralysis elicited by sleep interruption. Sleep, 15(3), 217–225. 10.1093/sleep/15.3.217

Tart, C. T. (1967). A second psychophysiological study of the out-of-the-body experiences in a selected subject. The Internal Journal of Parapsychology, 9(3), 251–258.

Tart, C. T. (1968). A Psychophysiological Study of Out-of-the-Body Experiences in a Selected Subject. The American Society for Psychical Research, 62, 3–27.

Terzaghi, M., Ratti, P. L., Manni, F., & Manni, R. (2012). Sleep paralysis in narcolepsy: More than just a motor dissociative phenomenon? Neurological Sciences: Official Journal of the Italian Neurological Society and of the Italian Society of Clinical Neurophysiology, 33(1), 169–172. 10.1007/s10072-011-0644-y

Timmermann, C., Roseman, L., Schartner, M., Milliere, R., Williams, L. T. J., Erritzoe, D., Muthukumaraswamy, S., Ashton, M., Bendrioua, A., Kaur, O., Turton, S., Nour, M. M., Day, C. M., Leech, R., Nutt, D. J., & Carhart-Harris, R. L. (2019). Neural correlates of the DMT experience assessed with multivariate EEG. Scientific Reports, 9(1), 16324. 10.1038/s41598-019-51974-4

Tressoldi, P. E., Pederzoli, L., Caini, P., Ferrini, A., Melloni, S., Prati, E., Richeldi, D., Richeldi, F., & Trabucco, A. (2015). Hypnotically Induced Out-of-Body Experience:How Many Bodies Are There? Unexpected Discoveries About the Subtle Body and Psychic Body. SAGE Open, 5(4), 2158244015615919. 10.1177/2158244015615919

Twemlow, S. W., Gabbard, G. O., & Jones, F. C. (1982). The out-of-body experience: A phenomenological typology based on questionnaire responses. The American Journal of Psychiatry, 139(4), 450–455. 10.1176/ajp.139.4.450

Valenzuela-Moguillansky, C., & Vásquez-Rosati, A. (2019). An Analysis Procedure for the Micro-Phenomenological Interview. Constructivist Foundations, 14(2), 123–145.

Varela, F. J. (1996). NEUROPHENOMENOLOGY. A Methodological Remedy for the Hard Problem. Journal of Consciousness Studies, 3(4), 330–349.

Vohryzek, J., Garcia-Guzmán, E., Kringelbach, M. L., Lopez-Sola, E., Timmermann, C., Roseman, L., Tagliazucchi, E., Ruffini, G., Carhart-Harris, R., Deco, G., & Sanz Perl, Y. (2025). Brain dynamics of classical psychedelics show paradoxical hierarchical flattening with increased complexity. bioRxiv Preprint. 10.1101/2024.12.21.629922

Weiler, M., Casseb, R. F., & Acunzo, D. (2025). Using Eye Movements to Signal the Onset of Self-Induced Out-of-Body Experiences (OBEs). Journal of Consciousness Studies, 32, 135–149. 10.53765/20512201.32.3.135

Zeev-Wolf, M., Dor-Ziderman, Y., Goldstein, A., Bonne, O., & Abramowitz, E. G. (2017). Oscillatory brain mechanisms of the hypnotically-induced out-of-body experience. Cortex; a Journal Devoted to the Study of the Nervous System and Behavior, 96, 19–30. 10.1016/j.cortex.2017.08.025

Zhang, X. S., Roy, R. J., & Jensen, E. W. (2001). EEG complexity as a measure of depth of anesthesia for patients. IEEE Transactions on Bio-Medical Engineering, 48(12), 1424–1433. 10.1109/10.966601

