## Supplementary for "Flattened brain hierarchy and increased EEG complexity during unusual bodily experiences and out-of-body experiences"

#### Supplementary Material Overview

This supplementary material provides detailed statistical analyses that complement the main manuscript. The results are organized into two key sections: **(1) Supplementary figures**, which include frequency-band statistics for irreversibility values and their corresponding sensitivity analyses; and **(2) Supplementary tables**, divided in two subsections:

**(2.1) Model comparison statistics** based on likelihood ratio tests, which details the iterative procedure used to select the most optimal generalized linear mixed model. It includes the formula for each candidate model and multiple model comparison metrics, including the Akaike

Information Criterion (AIC), the Bayesian Information Criterion (BIC), Log-likelihood (LogLik), and p-values from likelihood ratio tests. The preferred model selected at each step of the procedure is also reported.

**(2.2) Statistical parameter estimates** for each contrast and generalized linear mixed model, including false discovery rate (FDR)-adjusted p-values. Results are reported for irreversibility values and hierarchy measures and include for each case: (i) broadband comparisons, (ii) broadband sensitivity analyses, (iii) frequency-band comparisons, and (iv) frequency-band sensitivity analyses. This section also includes results for complexity measures, limited to global broadband comparisons.

### 1. Supplementary figures

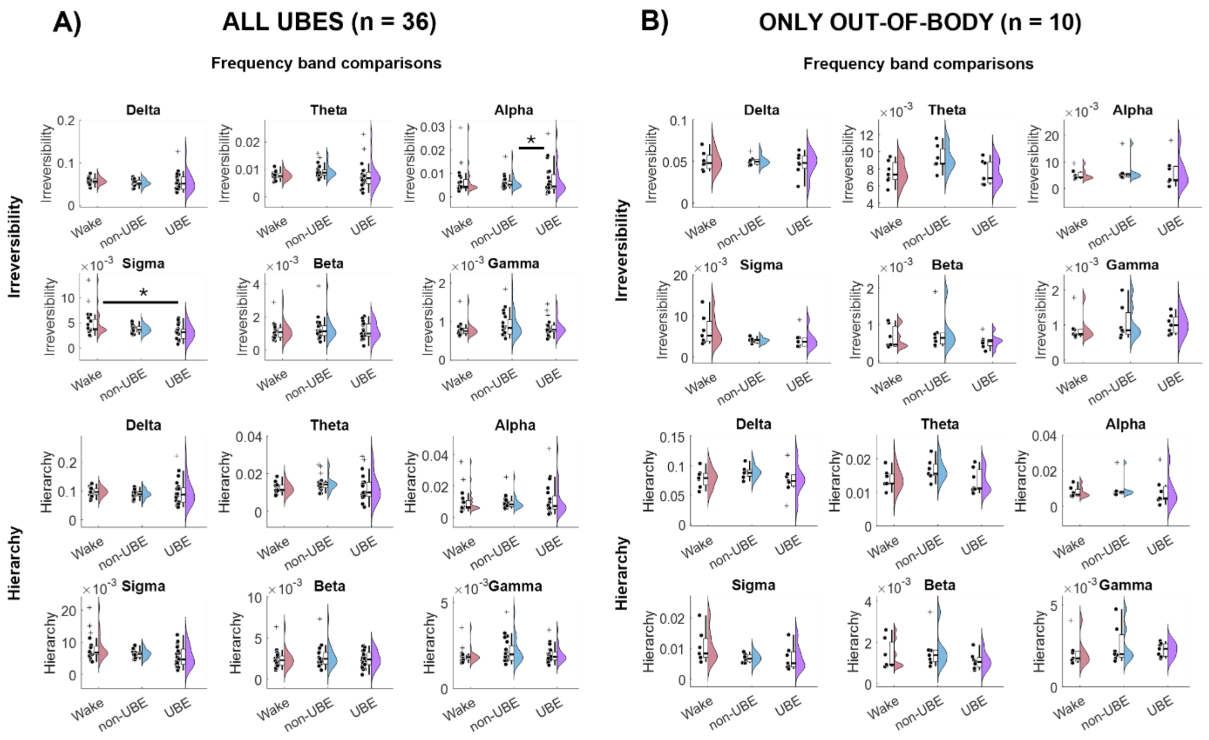

**Figure S1. EEG irreversibility across conditions within each frequency band. A)** Full UBE sample (n=36 UBEs, N=20 subjects). **B)** UBEs involving out-of-body experiences and/or elevated self-location experiences (n=10 UBEs, N=7 subjects). Dots indicate subject-level means for irreversibility (top) and hierarchy (bottom) within each frequency band: delta (1-4 Hz), theta (4-8 Hz), alpha (8-12 Hz), sigma (12-15 Hz), beta (15-30 Hz), and gamma (30-45 Hz) for visualization. Statistical comparisons with generalized linear mixed models were performed on individual trials following a within-subject design. Significance of model comparisons: *** $p < 0.001$, ** $p < 0.01$, * $p < 0.05$.

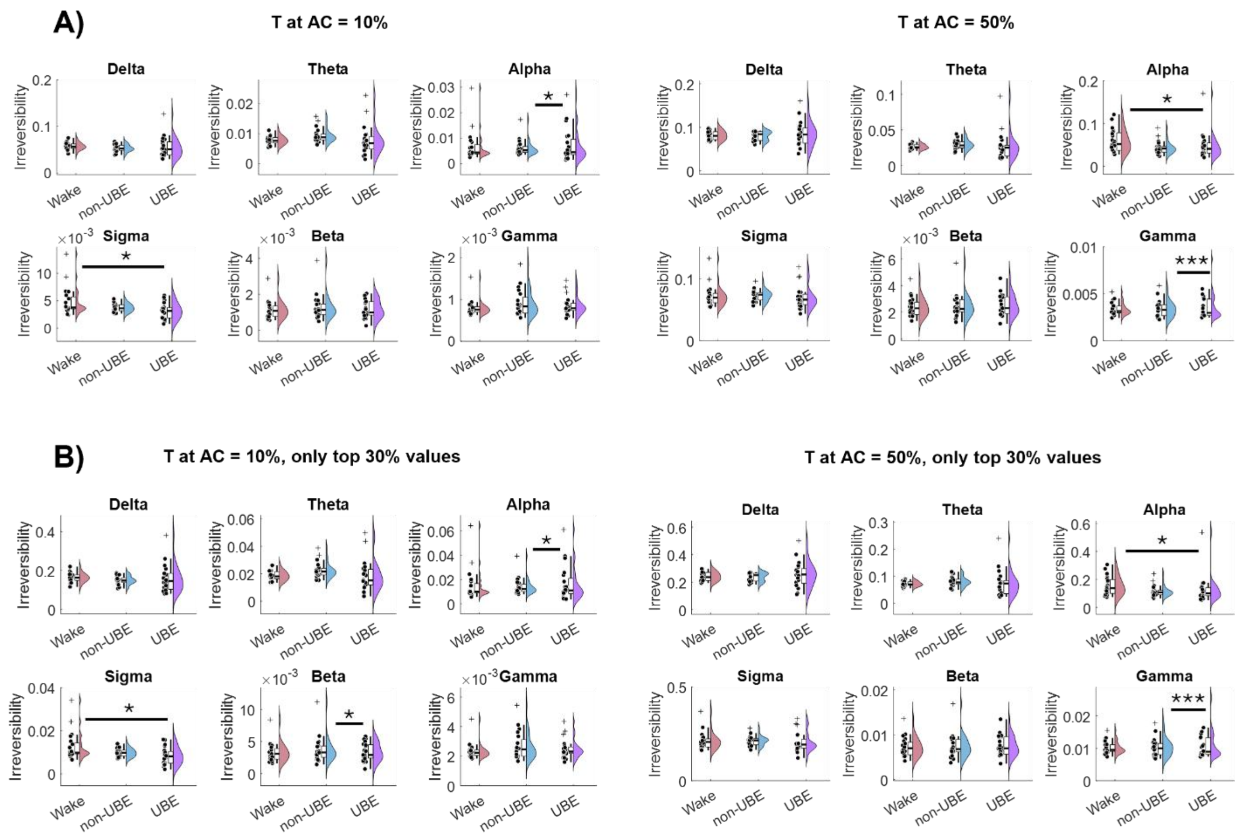

**Figure S2. Sensitivity analyses of EEG irreversibility within each frequency band. A)** Time lag T was tested at 10% and 50% of the maximum autocorrelation (AC). **B)** Only the top 30% irreversibility values were considered when T was set at either AC = 10% or 50%. Dots indicate subject-level means for irreversibility within each frequency band: delta (1-4 Hz), theta (4-8 Hz), alpha (8-12 Hz), sigma (12-15 Hz), beta (15-30 Hz), and gamma (30-45 Hz). Statistical comparisons with generalized linear mixed models were performed on individual trials following a within-subject design. Significance of model comparisons: *** $p < 0.001$, ** $p < 0.01$, * $p < 0.05$.

### 2. Supplementary tables

#### 2.1. Model comparison statistics – likelihood ratio tests

**Table S1: Likelihood ratio test results for "irreversibility" generalized linear mixed models**

| Step | Comparison | Model | Model formula | DF | AIC | BIC | LogLik | LRStat | deltaDF | pValue | Preferred model |
|---|---|---|---|---|---|---|---|---|---|---|---|
| **0** | glmm 0 vs. glmm 1 | glmm 0 | irreversibility ~ 1 | 2 | 26504 | 26492 | 13254 | 34.9080 | 2 | 0.0000 | glmm 1 |
| | | glmm 1 | irreversibility ~ 1 + Cond | 4 | 26535 | 26511 | 13271 | | | | |
| **1** | glmm 1 vs. glmm 2 | glmm 1 | irreversibility ~ 1 + Cond | 4 | 26535 | 26511 | 13271 | 562.0700 | 1 | 0.0000 | glmm 2 |
| | | glmm 2 | irreversibility ~ 1 + Cond + (1\|Subj) | 5 | 27095 | 27066 | 13553 | | | | |
| **2** | glmm 2 vs. glmm 3 | glmm 2 | irreversibility ~ 1 + Cond + (1\|Subj) | 5 | 27095 | 27066 | 13553 | 434.4700 | 0 | 0.0000 | glmm 3 |
| | | glmm 3 | irreversibility ~ 1 + Cond + (1\|Subj:Sleep) | 5 | 27530 | 27500 | 13770 | | | | |
| **3** | glmm 3 vs. glmm 4 | glmm 3 | irreversibility ~ 1 + Cond + (1\|Subj:Sleep) | 5 | 27530 | 27500 | 13770 | 7.2638 | 5 | 0.2018 | glmm 3 |
| | | glmm 4 | irreversibility ~ 1 + Cond + (Cond\|Subj:Sleep) | 10 | 27527 | 27468 | 13773 | | | | |
| **4** | glmm 3 vs. glmm 5 | glmm 3 | irreversibility ~ 1 + Cond + (1\|Subj:Sleep) | 5 | 27530 | 27500 | 13770 | 0.2695 | 1 | 0.6037 | glmm 3 |
| | | glmm 5 | irreversibility ~ 1 + Cond + (1\|Subj) + (1\|Subj:Sleep) | 6 | 27528 | 27492 | 13770 | | | | |
| **5** | glmm 3 vs. glmm 6 | glmm 3 | irreversibility ~ 1 + Cond + (1\|Subj:Sleep) | 5 | 27530 | 27500 | 13770 | 0.2695 | 2 | 0.8739 | glmm 3 |
| | | glmm 6 | irreversibility ~ 1 + Cond + (1\|Subj) + (1\|Sleep) + (1\|Subj:Sleep) | 7 | 27526 | 27484 | 13770 | | | | |
| **Final preferred model: glmm 3** | | | | | | | | | | | |

**Table S2: Likelihood ratio test results for "hierarchy" generalized linear mixed models**

| Step | Comparison | Model | Model formula | DF | AIC | BIC | LogLik | LRStat | deltaDF | pValue | Preferred model |
|---|---|---|---|---|---|---|---|---|---|---|---|
| **0** | glmm 0 vs. glmm 1 | glmm 0 | hierarchy ~ 1 | 2 | 23544 | 23532 | 11774 | 29.4320 | 2 | 0.0000 | glmm 1 |
| | | glmm 1 | hierarchy ~ 1 + Cond | 4 | 23569 | 23545 | 11788 | | | | |
| **1** | glmm 1 vs. glmm 2 | glmm 1 | hierarchy ~ 1 + Cond | 4 | 23569 | 23545 | 11788 | 488.9600 | 1 | 0.0000 | glmm 2 |
| | | glmm 2 | hierarchy ~ 1 + Cond + (1\|Subj) | 5 | 24056 | 24026 | 12033 | | | | |

| | | | | | | | | | | | |
|---|---|---|---|---|---|---|---|---|---|---|---|
| **2** | glmm 2 vs. glmm 3 | glmm 2 | hierarchy ~ 1 + Cond + (1\|Subj) | 5 | 24056 | 24026 | 12033 | 363.7800 | 0 | 0.0000 | glmm 3 |
| | | glmm 3 | hierarchy ~ 1 + Cond + (1\|Subj:Sleep) | 5 | 24420 | 24390 | 12215 | | | | |
| **3** | glmm 3 vs. glmm 4 | glmm 3 | hierarchy ~ 1 + Cond + (1\|Subj:Sleep) | 5 | 24420 | 24390 | 12215 | 8.6562 | 5 | 0.1236 | glmm 3 |
| | | glmm 4 | hierarchy ~ 1 + Cond + (Cond\|Subj:Sleep) | 10 | 24418 | 24359 | 12219 | | | | |
| **4** | glmm 3 vs. glmm 5 | glmm 3 | hierarchy ~ 1 + Cond + (1\|Subj:Sleep) | 5 | 24420 | 24390 | 12215 | 0.3733 | 1 | 0.5412 | glmm 3 |
| | | glmm 5 | hierarchy ~ 1 + Cond + (1\|Subj) + (1\|Subj:Sleep) | 6 | 24418 | 24383 | 12215 | | | | |
| **5** | glmm 3 vs. glmm 6 | glmm 3 | hierarchy ~ 1 + Cond + (1\|Subj:Sleep) | 5 | 24420 | 24390 | 12215 | 0.3733 | 2 | 0.8298 | glmm 3 |
| | | glmm 6 | hierarchy ~ 1 + Cond + (1\|Subj) + (1\|Sleep) + (1\|Subj:Sleep) | 7 | 24416 | 24375 | 12215 | | | | |
| **Final preferred model: glmm 3** | | | | | | | | | | | |

#### 2.2. Statistical parameter estimates - generalized linear mixed models

##### Irreversibility – Broadband comparisons

**Table S3: Broadband analysis – full UBE sample (T=10% of max autocorrelation)**

| | | | | | | CI 95% | | |
|---|---|---|---|---|---|---|---|---|
| **Contrast** | **Estimate** | **SE** | **tStat** | **DF** | **pvalue** | **lower** | **upper** | **FDR-adjusted p** |
| **'Intercept ascendent model'** | -6.0951 | 0.1249 | -48.8040 | 2690 | 0.0000 | -6.3400 | -5.8502 | - |
| **'Intercept descendent model'** | -6.1490 | 0.1460 | -42.1031 | 2690 | 0.0000 | -6.4354 | -5.8626 | - |
| **'Wake vs non-UBE'** | -0.0539 | 0.1922 | -0.2804 | 2690 | 0.7792 | -0.4307 | 0.3229 | 0.7792 |
| **'UBE vs non-UBE'** | -0.3929 | 0.0863 | -4.5525 | 2690 | 0.0000 | -0.5621 | -0.2237 | 0.0000 |
| **'UBE vs Wake'** | -0.3390 | 0.2017 | -1.6807 | 2690 | 0.0929 | -0.7345 | 0.0565 | 0.1394 |

**Table S4: Broadband analysis – out-of-body experiences (T=10% of max autocorrelation)**

| | | | | | | CI 95% | | |
|---|---|---|---|---|---|---|---|---|
| **Contrast** | **Estimate** | **SE** | **tStat** | **DF** | **pvalue** | **lower** | **upper** | **FDR-adjusted p** |
| **'Intercept ascendent model'** | -6.2857 | 0.1830 | -34.3490 | 870 | 0.0000 | -6.6448 | -5.9265 | - |
| **'Intercept descendent model'** | -6.0458 | 0.2021 | -29.9100 | 870 | 0.0000 | -6.4425 | -5.6491 | - |
| **'Wake vs non-UBE'** | 0.2399 | 0.2727 | 0.8797 | 870 | 0.3793 | -0.2953 | 0.7750 | 0.3793 |
| **'UBE vs non-UBE'** | -0.5131 | 0.1532 | -3.3479 | 870 | 0.0008 | -0.8138 | -0.2123 | 0.0025 |
| **'UBE vs Wake'** | -0.7529 | 0.2914 | -2.5838 | 870 | 0.0099 | -1.3249 | -0.1810 | 0.0149 |

##### Irreversibility – Broadband sensitivity analyses

**Table S5: Broadband sensitivity analysis – full UBE sample (T=0% of max autocorrelation)**

| | | | | | | CI 95% | | |
|---|---|---|---|---|---|---|---|---|
| **Contrast** | **Estimate** | **SE** | **tStat** | **DF** | **pvalue** | **lower** | **upper** | **FDR-adjusted p** |
| **'Intercept ascendent model'** | -6.1240 | 0.1145 | -53.4620 | 2690 | 0.0000 | -6.3486 | -5.8994 | - |
| **'Intercept descendent model'** | -6.1478 | 0.1333 | -46.1214 | 2690 | 0.0000 | -6.4092 | -5.8864 | - |
| **'Wake vs non-UBE'** | -0.0238 | 0.1758 | -0.1353 | 2690 | 0.8924 | -0.3684 | 0.3208 | 0.8924 |
| **'UBE vs non-UBE'** | -0.3496 | 0.0839 | -4.1653 | 2690 | 0.0000 | -0.5141 | -0.1850 | 0.0001 |
| **'UBE vs Wake'** | -0.3258 | 0.1858 | -1.7530 | 2690 | 0.0797 | -0.6901 | 0.0386 | 0.1196 |

**Table S6: Broadband sensitivity analysis – full UBE sample (T=20% of max autocorrelation)**

| | | | | | | CI 95% | | |
|---|---|---|---|---|---|---|---|---|
| **Contrast** | **Estimate** | **SE** | **tStat** | **DF** | **pvalue** | **lower** | **upper** | **FDR-adjusted p** |
| **'Intercept ascendent model'** | -6.1171 | 0.1183 | -51.7180 | 2690 | 0.0000 | -6.3490 | -5.8852 | - |

| | | | | | | | | |
|---|---|---|---|---|---|---|---|---|
| **'Intercept descendent model'** | -6.2275 | 0.1378 | -45.1805 | 2690 | 0.0000 | -6.4978 | -5.9573 | - |
| **'Wake vs non-UBE'** | -0.1104 | 0.1816 | -0.6081 | 2690 | 0.5432 | -0.4666 | 0.2457 | 0.5432 |
| **'UBE vs non-UBE'** | -0.4035 | 0.0821 | -4.9143 | 2690 | 0.0000 | -0.5644 | -0.2425 | 0.0000 |
| **'UBE vs Wake'** | -0.2930 | 0.1909 | -1.5348 | 2690 | 0.1249 | -0.6673 | 0.0813 | 0.1874 |

**Table S7: Broadband sensitivity analysis – full UBE sample (T=30% of max autocorrelation)**

| | | | | | | CI 95% | | |
|---|---|---|---|---|---|---|---|---|
| **Contrast** | **Estimate** | **SE** | **tStat** | **DF** | **pvalue** | **lower** | **upper** | **FDR-adjusted p** |
| **'Intercept ascendent model'** | -6.3145 | 0.1150 | -54.9028 | 2690 | 0.0000 | -6.5400 | -6.0889 | - |
| **'Intercept descendent model'** | -6.3849 | 0.1338 | -47.7040 | 2690 | 0.0000 | -6.6473 | -6.1224 | - |
| **'Wake vs non-UBE'** | -0.0704 | 0.1765 | -0.3991 | 2690 | 0.6899 | -0.4165 | 0.2756 | 0.6899 |
| **'UBE vs non-UBE'** | -0.3332 | 0.0789 | -4.2253 | 2690 | 0.0000 | -0.4879 | -0.1786 | 0.0001 |
| **'UBE vs Wake'** | -0.2628 | 0.1854 | -1.4175 | 2690 | 0.1564 | -0.6263 | 0.1007 | 0.2347 |

**Table S8: Broadband sensitivity analysis – full UBE sample (T=40% of max autocorrelation)**

| | | | | | | CI 95% | | |
|---|---|---|---|---|---|---|---|---|
| **Contrast** | **Estimate** | **SE** | **tStat** | **DF** | **pvalue** | **lower** | **upper** | **FDR-adjusted p** |
| **'Intercept ascendent model'** | -6.3283 | 0.1149 | -55.0579 | 2690 | 0.0000 | -6.5537 | -6.1029 | - |
| **'Intercept descendent model'** | -6.4605 | 0.1341 | -48.1629 | 2690 | 0.0000 | -6.7236 | -6.1975 | - |
| **'Wake vs non-UBE'** | -0.1322 | 0.1766 | -0.7486 | 2690 | 0.4542 | -0.4786 | 0.2141 | 0.4542 |
| **'UBE vs non-UBE'** | -0.4262 | 0.0793 | -5.3765 | 2690 | 0.0000 | -0.5817 | -0.2708 | 0.0000 |
| **'UBE vs Wake'** | -0.2940 | 0.1855 | -1.5849 | 2690 | 0.1131 | -0.6578 | 0.0697 | 0.1696 |

**Table S9: Broadband sensitivity analysis – full UBE sample (T=50% of max autocorrelation)**

| | | | | | | CI 95% | | |
|---|---|---|---|---|---|---|---|---|
| **Contrast** | **Estimate** | **SE** | **tStat** | **DF** | **pvalue** | **lower** | **upper** | **FDR-adjusted p** |
| **'Intercept ascendent model'** | -6.3146 | 0.1341 | -47.1056 | 2690 | 0.0000 | -6.5775 | -6.0518 | - |
| **'Intercept descendent model'** | -6.0955 | 0.1576 | -38.6820 | 2690 | 0.0000 | -6.4045 | -5.7865 | - |
| **'Wake vs non-UBE'** | 0.2191 | 0.2069 | 1.0591 | 2690 | 0.2897 | -0.1866 | 0.6248 | 0.2897 |
| **'UBE vs non-UBE'** | -0.4148 | 0.0840 | -4.9382 | 2690 | 0.0000 | -0.5795 | -0.2501 | 0.0000 |
| **'UBE vs Wake'** | -0.6339 | 0.2151 | -2.9464 | 2690 | 0.0032 | -1.0558 | -0.2120 | 0.0049 |

**Table S10: Broadband sensitivity analysis – full UBE sample (T=10% of max autocorrelation, using only 20% highest irreversibility values)**

| | | | | | | CI 95% | | |
|---|---|---|---|---|---|---|---|---|
| **Contrast** | **Estimate** | **SE** | **tStat** | **DF** | **pvalue** | **lower** | **upper** | **FDR-adjusted p** |
| **'Intercept ascendent model'** | -4.7761 | 0.1233 | -38.7267 | 2690 | 0.0000 | -5.0180 | -4.5343 | - |
| **'Intercept descendent model'** | -4.8464 | 0.1439 | -33.6794 | 2690 | 0.0000 | -5.1285 | -4.5642 | - |
| **'Wake vs non-UBE'** | -0.0702 | 0.1895 | -0.3706 | 2690 | 0.7109 | -0.4419 | 0.3014 | 0.7109 |
| **'UBE vs non-UBE'** | -0.3829 | 0.0886 | -4.3197 | 2690 | 0.0000 | -0.5567 | -0.2091 | 0.0000 |
| **'UBE vs Wake'** | -0.3127 | 0.1998 | -1.5650 | 2690 | 0.1177 | -0.7044 | 0.0791 | 0.1765 |

**Table S11: Broadband sensitivity analysis – full UBE sample (T=50% of max autocorrelation, using only 20% highest irreversibility values)**

| | | | | | | CI 95% | | |
|---|---|---|---|---|---|---|---|---|
| **Contrast** | **Estimate** | **SE** | **tStat** | **DF** | **pvalue** | **lower** | **upper** | **FDR-adjusted p** |
| **'Intercept ascendent model'** | -4.9330 | 0.1312 | -37.5958 | 2690 | 0.0000 | -5.1903 | -4.6757 | - |
| **'Intercept descendent model'** | -4.7405 | 0.1538 | -30.8172 | 2690 | 0.0000 | -5.0422 | -4.4389 | - |
| **'Wake vs non-UBE'** | 0.1925 | 0.2022 | 0.9520 | 2690 | 0.3412 | -0.2040 | 0.5889 | 0.3412 |
| **'UBE vs non-UBE'** | -0.4178 | 0.0854 | -4.8906 | 2690 | 0.0000 | -0.5853 | -0.2503 | 0.0000 |
| **'UBE vs Wake'** | -0.6103 | 0.2110 | -2.8918 | 2690 | 0.0039 | -1.0241 | -0.1965 | 0.0058 |

**Table S12: Broadband sensitivity analysis – REM sleep group (T=10% of max autocorrelation)**

| | | | | | | CI 95% | | |
|---|---|---|---|---|---|---|---|---|
| **Contrast** | **Estimate** | **SE** | **tStat** | **DF** | **pvalue** | **lower** | **upper** | **FDR-adjusted p** |
| **'Intercept ascendent model'** | -5.9470 | 0.1618 | -36.7624 | 605 | 0.0000 | -6.2647 | -5.6293 | - |
| **'Intercept descendent model'** | -5.8187 | 0.1627 | -35.7571 | 605 | 0.0000 | -6.1383 | -5.4991 | - |
| **'Wake vs non-UBE'** | 0.1283 | 0.0937 | 1.3692 | 605 | 0.1715 | -0.0557 | 0.3123 | 0.1715 |
| **'UBE vs non-UBE'** | -0.5370 | 0.2118 | -2.5356 | 605 | 0.0115 | -0.9529 | -0.1211 | 0.0172 |
| **'UBE vs Wake'** | -0.6653 | 0.2110 | -3.1529 | 605 | 0.0017 | -1.0797 | -0.2509 | 0.0051 |

**Table S13: Broadband sensitivity analysis – Light sleep group (T=10% of max autocorrelation)**

| | | | | | | CI 95% | | |
|---|---|---|---|---|---|---|---|---|
| **Contrast** | **Estimate** | **SE** | **tStat** | **DF** | **pvalue** | **lower** | **upper** | **FDR-adjusted p** |
| **'Intercept ascendent model'** | -5.5120 | 0.0647 | -85.1593 | 625 | 0.0000 | -5.6391 | -5.3848 | - |
| **'Intercept descendent model'** | -6.4884 | 0.0655 | -99.0693 | 625 | 0.0000 | -6.6170 | -6.3598 | - |
| **'Wake vs non-UBE'** | -0.9765 | 0.0921 | -10.6046 | 625 | 0.0000 | -1.1573 | -0.7956 | 0.0000 |
| **'UBE vs non-UBE'** | -0.3874 | 0.2002 | -1.9346 | 625 | 0.0535 | -0.7806 | 0.0058 | 0.0535 |
| **'UBE vs Wake'** | 0.5891 | 0.2005 | 2.9381 | 625 | 0.0034 | 0.1954 | 0.9828 | 0.0051 |

**Table S14: Broadband sensitivity analysis – Meditation group (T=10% of max autocorrelation)**

| | | | | | | CI 95% | | |
|---|---|---|---|---|---|---|---|---|
| **Contrast** | **Estimate** | **SE** | **tStat** | **DF** | **pvalue** | **lower** | **upper** | **FDR-adjusted p** |
| **'Intercept ascendent model'** | -6.4389 | 0.1572 | -40.9672 | 873 | 0.0000 | -6.7473 | -6.1304 | - |
| **'Intercept descendent model'** | -6.3245 | 0.1573 | -40.1950 | 873 | 0.0000 | -6.6333 | -6.0157 | - |
| **'Wake vs non-UBE'** | 0.1144 | 0.0678 | 1.6873 | 873 | 0.0919 | -0.0187 | 0.2474 | 0.0919 |
| **'UBE vs non-UBE'** | -0.6599 | 0.1270 | -5.1963 | 873 | 0.0000 | -0.9091 | -0.4106 | 0.0000 |
| **'UBE vs Wake'** | -0.7743 | 0.1266 | -6.1178 | 873 | 0.0000 | -1.0227 | -0.5259 | 0.0000 |

**Table S15: Broadband sensitivity analysis – Sleep arousal group (T=10% of max autocorrelation)**

| | | | | | | CI 95% |
|---|---|---|---|---|---|---|

| Contrast | Estimate | SE | tStat | DF | pvalue | lower | upper | FDR-adjusted p |
|---|---|---|---|---|---|---|---|---|
| 'Intercept ascendent model' | -5.7828 | 0.1740 | -33.2367 | 864 | 0.0000 | -6.1243 | -5.4413 | - |
| 'Intercept descendent model' | -6.2990 | 0.1743 | -36.1290 | 864 | 0.0000 | -6.6412 | -5.9568 | - |
| 'Wake vs non-UBE' | -0.5162 | 0.0784 | -6.5805 | 864 | 0.0000 | -0.6701 | -0.3622 | 0.0000 |
| 'UBE vs non-UBE' | -0.5352 | 0.1633 | -3.2769 | 864 | 0.0011 | -0.8558 | -0.2146 | 0.0016 |
| 'UBE vs Wake' | -0.0191 | 0.1577 | -0.1209 | 864 | 0.9038 | -0.3286 | 0.2905 | 0.9038 |

### Irreversibility – Frequency band comparisons

**Table S16: Frequency band analysis – full UBE sample (T=10% of max autocorrelation)**

| | | | | | | CI 95% | | |
|---|---|---|---|---|---|---|---|---|
| **Contrast** | **Estimate** | **SE** | **tStat** | **DF** | **pvalue** | **lower** | **upper** | **FDR-adjusted p** |
| **DELTA BAND** | | | | | | | | |
| Intercept ascendent model' | -2.9299 | 0.0289 | -101.3645 | 2690 | 0.0000 | -2.9866 | -2.8732 | - |
| 'Intercept descendent model' | -2.8601 | 0.0314 | -91.1094 | 2690 | 0.0000 | -2.9217 | -2.7986 | - |
| 'Wake vs non-UBE' | 0.0697 | 0.0427 | 1.6342 | 2690 | 0.1023 | -0.0139 | 0.1534 | 0.2604 |
| 'UBE vs non-UBE' | 0.0374 | 0.0663 | 0.5647 | 2690 | 0.5724 | -0.0925 | 0.1674 | 0.6868 |
| 'UBE vs Wake' | -0.0323 | 0.0711 | -0.4542 | 2690 | 0.6497 | -0.1718 | 0.1072 | 0.7309 |
| **THETA BAND** | | | | | | | | |
| Intercept ascendent model' | -4.7406 | 0.0487 | -97.3618 | 2690 | 0.0000 | -4.8360 | -4.6451 | - |
| 'Intercept descendent model' | -4.8656 | 0.0531 | -91.6265 | 2690 | 0.0000 | -4.9698 | -4.7615 | - |
| 'Wake vs non-UBE' | -0.1251 | 0.0720 | -1.7361 | 2690 | 0.0827 | -0.2664 | 0.0162 | 0.2480 |
| 'UBE vs non-UBE' | -0.1339 | 0.0851 | -1.5735 | 2690 | 0.1157 | -0.3008 | 0.0330 | 0.2604 |
| 'UBE vs Wake' | -0.0088 | 0.1017 | -0.0867 | 2690 | 0.9309 | -0.2083 | 0.1906 | 0.9309 |
| **ALPHA BAND** | | | | | | | | |
| 'Intercept ascendent model' | -5.0226 | 0.1007 | -49.8697 | 2690 | 0.0000 | -5.2200 | -4.8251 | - |
| 'Intercept descendent model' | -5.1985 | 0.1155 | -45.0068 | 2690 | 0.0000 | -5.4250 | -4.9720 | - |
| 'Wake vs non-UBE' | -0.1760 | 0.1532 | -1.1482 | 2690 | 0.2510 | -0.4764 | 0.1245 | 0.5020 |
| 'UBE vs non-UBE' | -0.2829 | 0.0995 | -2.8438 | 2690 | 0.0045 | -0.4780 | -0.0878 | 0.0412 |
| 'UBE vs Wake' | -0.1070 | 0.1700 | -0.6293 | 2690 | 0.5292 | -0.4403 | 0.2263 | 0.6804 |
| **SIGMA BAND** | | | | | | | | |
| 'Intercept ascendent model' | -5.6148 | 0.0657 | -85.4211 | 2690 | 0.0000 | -5.7437 | -5.4859 | - |
| 'Intercept descendent model' | -5.4233 | 0.0733 | -74.0281 | 2690 | 0.0000 | -5.5670 | -5.2797 | - |
| 'Wake vs non-UBE' | 0.1915 | 0.0984 | 1.9454 | 2690 | 0.0518 | -0.0015 | 0.3845 | 0.2332 |
| 'UBE vs non-UBE' | -0.1468 | 0.0841 | -1.7454 | 2690 | 0.0810 | -0.3118 | 0.0181 | 0.2480 |
| 'UBE vs Wake' | -0.3383 | 0.1192 | -2.8380 | 2690 | 0.0046 | -0.5720 | -0.1046 | 0.0412 |
| **BETA BAND** | | | | | | | | |
| 'Intercept ascendent model' | -6.7373 | 0.0810 | -83.2196 | 2690 | 0.0000 | -6.8960 | -6.5785 | - |
| 'Intercept descendent model' | -6.8549 | 0.0943 | -72.7157 | 2690 | 0.0000 | -7.0398 | -6.6701 | - |
| 'Wake vs non-UBE' | -0.1177 | 0.1243 | -0.9470 | 2690 | 0.3437 | -0.3613 | 0.1260 | 0.5625 |
| 'UBE vs non-UBE' | -0.1544 | 0.0611 | -2.5257 | 2690 | 0.0116 | -0.2742 | -0.0345 | 0.0696 |

| | | | | | | | | |
|---|---|---|---|---|---|---|---|---|
| 'UBE vs Wake' | -0.0367 | 0.1317 | -0.2786 | 2690 | 0.7806 | -0.2950 | 0.2216 | 0.8265 |
| **GAMMA BAND** | | | | | | | | |
| 'Intercept ascendent model' | -7.1042 | 0.0485 | -146.5381 | 2690 | 0.0000 | -7.1993 | -7.0092 | - |
| 'Intercept descendent model' | -7.1543 | 0.0561 | -127.5030 | 2690 | 0.0000 | -7.2643 | -7.0443 | - |
| 'Wake vs non-UBE' | -0.0501 | 0.0742 | -0.6754 | 2690 | 0.4995 | -0.1955 | 0.0953 | 0.6804 |
| 'UBE vs non-UBE' | 0.0340 | 0.0385 | 0.8828 | 2690 | 0.3774 | -0.0415 | 0.1095 | 0.5661 |
| 'UBE vs Wake' | 0.0841 | 0.0793 | 1.0607 | 2690 | 0.2889 | -0.0714 | 0.2395 | 0.5200 |

**Table S17: Frequency band analysis – out-of-body experiences (T=10% of max autocorrelation)**

| | | | | | | CI 95% | | |
|---|---|---|---|---|---|---|---|---|
| **Contrast** | **Estimate** | **SE** | **tStat** | **DF** | **pvalue** | **lower** | **upper** | **FDR-adjusted p** |
| **DELTA BAND** | | | | | | | | |
| 'Intercept ascendent model' | -2.9792 | 0.0564 | -52.7942 | 870 | 0.0000 | -3.0899 | -2.8684 | - |
| 'Intercept descendent model' | -3.0045 | 0.0574 | -52.3816 | 870 | 0.0000 | -3.1170 | -2.8919 | - |
| 'Wake vs non-UBE' | -0.0253 | 0.0805 | -0.3143 | 870 | 0.7534 | -0.1832 | 0.1326 | 0.8827 |
| 'UBE vs non-UBE' | -0.0526 | 0.1201 | -0.4381 | 870 | 0.6614 | -0.2882 | 0.1830 | 0.8827 |
| 'UBE vs Wake' | -0.0273 | 0.1300 | -0.2100 | 870 | 0.8337 | -0.2825 | 0.2278 | 0.8827 |
| **THETA BAND** | | | | | | | | |
| 'Intercept ascendent model' | -4.6993 | 0.0584 | -80.5175 | 870 | 0.0000 | -4.8139 | -4.5848 | - |
| 'Intercept descendent model' | -4.8888 | 0.0593 | -82.4286 | 870 | 0.0000 | -5.0052 | -4.7724 | - |
| 'Wake vs non-UBE' | -0.1895 | 0.0832 | -2.2769 | 870 | 0.0230 | -0.3528 | -0.0261 | 0.2404 |
| 'UBE vs non-UBE' | -0.1601 | 0.1294 | -1.2377 | 870 | 0.2162 | -0.4140 | 0.0938 | 0.4864 |
| 'UBE vs Wake' | 0.0294 | 0.1394 | 0.2105 | 870 | 0.8333 | -0.2443 | 0.3030 | 0.8827 |
| **ALPHA BAND** | | | | | | | | |
| 'Intercept ascendent model' | -4.9968 | 0.1616 | -30.9218 | 870 | 0.0000 | -5.3140 | -4.6796 | - |
| 'Intercept descendent model' | -5.3069 | 0.1737 | -30.5526 | 870 | 0.0000 | -5.6478 | -4.9660 | - |
| 'Wake vs non-UBE' | -0.3101 | 0.2372 | -1.3071 | 870 | 0.1915 | -0.7757 | 0.1555 | 0.4864 |
| 'UBE vs non-UBE' | -0.3912 | 0.1873 | -2.0890 | 870 | 0.0370 | -0.7587 | -0.0236 | 0.2404 |
| 'UBE vs Wake' | -0.0811 | 0.2721 | -0.2980 | 870 | 0.7658 | -0.6151 | 0.4529 | 0.8827 |
| **SIGMA BAND** | | | | | | | | |
| 'Intercept ascendent model' | -5.4811 | 0.1284 | -42.6729 | 870 | 0.0000 | -5.7332 | -5.2290 | - |
| 'Intercept descendent model' | -5.1538 | 0.1355 | -38.0323 | 870 | 0.0000 | -5.4197 | -4.8878 | - |
| 'Wake vs non-UBE' | 0.3273 | 0.1867 | 1.7529 | 870 | 0.0800 | -0.0392 | 0.6937 | 0.2879 |
| 'UBE vs non-UBE' | -0.1055 | 0.1640 | -0.6434 | 870 | 0.5201 | -0.4274 | 0.2164 | 0.8827 |
| 'UBE vs Wake' | -0.4328 | 0.2238 | -1.9341 | 870 | 0.0534 | -0.8720 | 0.0064 | 0.2404 |
| **BETA BAND** | | | | | | | | |
| 'Intercept ascendent model' | -7.3501 | 0.1443 | -50.9300 | 870 | 0.0000 | -7.6334 | -7.0669 | - |
| 'Intercept descendent model' | -7.4409 | 0.1584 | -46.9900 | 870 | 0.0000 | -7.7517 | -7.1301 | - |
| 'Wake vs non-UBE' | -0.0907 | 0.2142 | -0.4235 | 870 | 0.6720 | -0.5112 | 0.3298 | 0.8827 |
| 'UBE vs non-UBE' | -0.0698 | 0.1241 | -0.5627 | 870 | 0.5738 | -0.3134 | 0.1738 | 0.8827 |
| 'UBE vs Wake' | 0.0209 | 0.2303 | 0.0908 | 870 | 0.9277 | -0.4311 | 0.4729 | 0.9277 |
| **GAMMA BAND** | | | | | | | | |

| | | | | | | | | |
|---|---|---|---|---|---|---|---|---|
| **'Intercept ascendent model'** | -6.9843 | 0.1130 | -61.8255 | 870 | 0.0000 | -7.2060 | -6.7626 | - |
| **'Intercept descendent model'** | -7.0625 | 0.1259 | -56.0923 | 870 | 0.0000 | -7.3097 | -6.8154 | - |
| **'Wake vs non-UBE'** | -0.0782 | 0.1692 | -0.4625 | 870 | 0.6438 | -0.4102 | 0.2538 | 0.8827 |
| **'UBE vs non-UBE'** | 0.1760 | 0.0859 | 2.0490 | 870 | 0.0408 | 0.0074 | 0.3446 | 0.2404 |
| **'UBE vs Wake'** | 0.2542 | 0.1782 | 1.4265 | 870 | 0.1541 | -0.0956 | 0.6040 | 0.4623 |

### Irreversibility – Frequency band sensitivity analyses

**Table S17: Frequency band sensitivity analysis – full UBE sample (T=50% of max autocorrelation)**

| | | | | | | **CI 95%** | | |
|---|---|---|---|---|---|---|---|---|
| **Contrast** | **Estimate** | **SE** | **tStat** | **DF** | **pvalue** | **lower** | **upper** | **FDR-adjusted p** |
| | | | | **DELTA BAND** | | | | |
| **'Intercept ascendent model'** | -2.5203 | 0.0251 | -100.3631 | 2690 | 0.0000 | -2.5696 | -2.4711 | - |
| **'Intercept descendent model'** | -2.5141 | 0.0273 | -92.2336 | 2690 | 0.0000 | -2.5675 | -2.4606 | - |
| **'Wake vs non-UBE'** | 0.0062 | 0.0371 | 0.1685 | 2690 | 0.8662 | -0.0664 | 0.0789 | 0.9149 |
| **'UBE vs non-UBE'** | 0.0390 | 0.0593 | 0.6578 | 2690 | 0.5108 | -0.0773 | 0.1552 | 0.8700 |
| **'UBE vs Wake'** | 0.0328 | 0.0631 | 0.5191 | 2690 | 0.6037 | -0.0910 | 0.1565 | 0.9056 |
| | | | | **THETA BAND** | | | | |
| **'Intercept ascendent model'** | -3.5602 | 0.0402 | -88.4911 | 2690 | 0.0000 | -3.6391 | -3.4814 | - |
| **'Intercept descendent model'** | -3.6332 | 0.0438 | -83.0404 | 2690 | 0.0000 | -3.7190 | -3.5474 | - |
| **'Wake vs non-UBE'** | -0.0729 | 0.0594 | -1.2270 | 2690 | 0.2199 | -0.1895 | 0.0436 | 0.4949 |
| **'UBE vs non-UBE'** | -0.1094 | 0.0757 | -1.4456 | 2690 | 0.1484 | -0.2578 | 0.0390 | 0.4949 |
| **'UBE vs Wake'** | -0.0365 | 0.0877 | -0.4162 | 2690 | 0.6773 | -0.2084 | 0.1354 | 0.9149 |
| | | | | **ALPHA BAND** | | | | |
| **'Intercept ascendent model'** | -3.1002 | 0.0722 | -42.9283 | 2690 | 0.0000 | -3.2418 | -2.9586 | - |
| **'Intercept descendent model'** | -2.8993 | 0.0817 | -35.4825 | 2690 | 0.0000 | -3.0595 | -2.7390 | - |
| **'Wake vs non-UBE'** | 0.2009 | 0.1091 | 1.8427 | 2690 | 0.0655 | -0.0129 | 0.4148 | 0.2947 |
| **'UBE vs non-UBE'** | -0.1651 | 0.0816 | -2.0239 | 2690 | 0.0431 | -0.3250 | -0.0051 | 0.2585 |
| **'UBE vs Wake'** | -0.3660 | 0.1256 | -2.9144 | 2690 | 0.0036 | -0.6122 | -0.1197 | 0.0323 |
| | | | | **SIGMA BAND** | | | | |
| **'Intercept ascendent model'** | -2.6176 | 0.0338 | -77.4375 | 2690 | 0.0000 | -2.6838 | -2.5513 | - |
| **'Intercept descendent model'** | -2.6489 | 0.0369 | -71.7316 | 2690 | 0.0000 | -2.7213 | -2.5765 | - |
| **'Wake vs non-UBE'** | -0.0313 | 0.0501 | -0.6255 | 2690 | 0.5317 | -0.1295 | 0.0668 | 0.8700 |
| **'UBE vs non-UBE'** | -0.0521 | 0.0628 | -0.8302 | 2690 | 0.4065 | -0.1752 | 0.0710 | 0.8130 |
| **'UBE vs Wake'** | -0.0208 | 0.0726 | -0.2869 | 2690 | 0.7742 | -0.1631 | 0.1214 | 0.9149 |
| | | | | **BETA BAND** | | | | |
| **'Intercept ascendent model'** | -6.0462 | 0.0590 | -102.4592 | 2690 | 0.0000 | -6.1619 | -5.9305 | - |
| **'Intercept descendent model'** | -6.0690 | 0.0675 | -89.8787 | 2690 | 0.0000 | -6.2014 | -5.9366 | - |
| **'Wake vs non-UBE'** | -0.0228 | 0.0897 | -0.2546 | 2690 | 0.7991 | -0.1987 | 0.1530 | 0.9149 |
| **'UBE vs non-UBE'** | -0.0059 | 0.0554 | -0.1068 | 2690 | 0.9149 | -0.1145 | 0.1027 | 0.9149 |
| **'UBE vs Wake'** | 0.0169 | 0.0987 | 0.1713 | 2690 | 0.8640 | -0.1767 | 0.2105 | 0.9149 |

| GAMMA BAND | | | | | | | | |
|---|---|---|---|---|---|---|---|---|
| **'Intercept ascendent model'** | -5.8009 | 0.0466 | -124.5740 | 2690 | 0.0000 | -5.8922 | -5.7095 | - |
| **'Intercept descendent model'** | -5.7049 | 0.0540 | -105.6396 | 2690 | 0.0000 | -5.8108 | -5.5990 | - |
| **'Wake vs non-UBE'** | 0.0960 | 0.0713 | 1.3458 | 2690 | 0.1785 | -0.0439 | 0.2358 | 0.4949 |
| **'UBE vs non-UBE'** | 0.1929 | 0.0357 | 5.4073 | 2690 | 0.0000 | 0.1229 | 0.2628 | 0.0000 |
| **'UBE vs Wake'** | 0.0969 | 0.0758 | 1.2778 | 2690 | 0.2014 | -0.0518 | 0.2456 | 0.4949 |

**Table S18: Frequency band sensitivity analysis – full UBE sample (T=10% of max autocorrelation, using only 30% highest irreversibility values)**

| | | | | | | CI 95% | | |
|---|---|---|---|---|---|---|---|---|
| **Contrast** | **Estimate** | **SE** | **tStat** | **DF** | **pvalue** | **lower** | **upper** | **FDR-adjusted p** |
| **DELTA BAND** | | | | | | | | |
| **'Intercept ascendent model'** | -1.8983 | 0.0300 | -63.1728 | 2690 | 0.0000 | -1.9572 | -1.8393 | - |
| **'Intercept descendent model'** | -1.8201 | 0.0326 | -55.7612 | 2690 | 0.0000 | -1.8842 | -1.7561 | - |
| **'Wake vs non-UBE'** | 0.0781 | 0.0444 | 1.7606 | 2690 | 0.0784 | -0.0089 | 0.1651 | 0.2176 |
| **'UBE vs non-UBE'** | 0.0529 | 0.0676 | 0.7824 | 2690 | 0.4341 | -0.0796 | 0.1853 | 0.6278 |
| **'UBE vs Wake'** | -0.0253 | 0.0730 | -0.3462 | 2690 | 0.7292 | -0.1683 | 0.1178 | 0.7721 |
| **THETA BAND** | | | | | | | | |
| **'Intercept ascendent model'** | -3.8559 | 0.0493 | -78.1374 | 2690 | 0.0000 | -3.9526 | -3.7591 | - |
| **'Intercept descendent model'** | -4.0015 | 0.0538 | -74.3729 | 2690 | 0.0000 | -4.1070 | -3.8960 | - |
| **'Wake vs non-UBE'** | -0.1456 | 0.0730 | -1.9946 | 2690 | 0.0462 | -0.2888 | -0.0025 | 0.1796 |
| **'UBE vs non-UBE'** | -0.1490 | 0.0866 | -1.7201 | 2690 | 0.0855 | -0.3188 | 0.0208 | 0.2176 |
| **'UBE vs Wake'** | -0.0033 | 0.1033 | -0.0324 | 2690 | 0.9742 | -0.2060 | 0.1993 | 0.9742 |
| **ALPHA BAND** | | | | | | | | |
| **'Intercept ascendent model'** | -4.1835 | 0.0977 | -42.8112 | 2690 | 0.0000 | -4.3751 | -3.9919 | - |
| **'Intercept descendent model'** | -4.3611 | 0.1117 | -39.0461 | 2690 | 0.0000 | -4.5801 | -4.1421 | - |
| **'Wake vs non-UBE'** | -0.1776 | 0.1484 | -1.1965 | 2690 | 0.2316 | -0.4686 | 0.1134 | 0.4632 |
| **'UBE vs non-UBE'** | -0.2836 | 0.1003 | -2.8282 | 2690 | 0.0047 | -0.4803 | -0.0870 | 0.0318 |
| **'UBE vs Wake'** | -0.1061 | 0.1661 | -0.6385 | 2690 | 0.5232 | -0.4319 | 0.2197 | 0.6278 |
| **SIGMA BAND** | | | | | | | | |
| **'Intercept ascendent model'** | -4.6506 | 0.0650 | -71.5888 | 2690 | 0.0000 | -4.7780 | -4.5233 | - |
| **'Intercept descendent model'** | -4.4600 | 0.0723 | -61.6960 | 2690 | 0.0000 | -4.6017 | -4.3182 | - |
| **'Wake vs non-UBE'** | 0.1907 | 0.0972 | 1.9617 | 2690 | 0.0499 | 0.0001 | 0.3812 | 0.1796 |
| **'UBE vs non-UBE'** | -0.1413 | 0.0851 | -1.6615 | 2690 | 0.0967 | -0.3081 | 0.0255 | 0.2176 |
| **'UBE vs Wake'** | -0.3320 | 0.1188 | -2.7953 | 2690 | 0.0052 | -0.5649 | -0.0991 | 0.0318 |
| **BETA BAND** | | | | | | | | |
| **'Intercept ascendent model'** | -5.6696 | 0.0813 | -69.7067 | 2690 | 0.0000 | -5.8291 | -5.5102 | - |
| **'Intercept descendent model'** | -5.7884 | 0.0946 | -61.1615 | 2690 | 0.0000 | -5.9739 | -5.6028 | - |
| **'Wake vs non-UBE'** | -0.1187 | 0.1248 | -0.9513 | 2690 | 0.3415 | -0.3634 | 0.1260 | 0.5588 |
| **'UBE vs non-UBE'** | -0.1728 | 0.0619 | -2.7903 | 2690 | 0.0053 | -0.2942 | -0.0514 | 0.0318 |
| **'UBE vs Wake'** | -0.0541 | 0.1325 | -0.4081 | 2690 | 0.6833 | -0.3138 | 0.2057 | 0.7687 |
| **GAMMA BAND** | | | | | | | | |

| | | | | | | | | |
|---|---|---|---|---|---|---|---|---|
| 'Intercept ascendent model' | -6.0209 | 0.0485 | -124.1508 | 2690 | 0.0000 | -6.1160 | -5.9258 | - |
| 'Intercept descendent model' | -6.0759 | 0.0561 | -108.2758 | 2690 | 0.0000 | -6.1859 | -5.9658 | - |
| 'Wake vs non-UBE' | -0.0550 | 0.0742 | -0.7410 | 2690 | 0.4588 | -0.2004 | 0.0905 | 0.6278 |
| 'UBE vs non-UBE' | 0.0263 | 0.0386 | 0.6816 | 2690 | 0.4955 | -0.0494 | 0.1021 | 0.6278 |
| 'UBE vs Wake' | 0.0813 | 0.0793 | 1.0248 | 2690 | 0.3055 | -0.0742 | 0.2368 | 0.5500 |

**Table S19: Frequency band sensitivity analysis – full UBE sample (T=50% of max autocorrelation, using only 30% highest irreversibility values)**

| | | | | | | CI 95% | | |
|---|---|---|---|---|---|---|---|---|
| **Contrast** | **Estimate** | **SE** | **tStat** | **DF** | **pvalue** | **lower** | **upper** | **FDR-adjusted p** |
| **DELTA BAND** | | | | | | | | |
| 'Intercept ascendent model' | -1.4343 | 0.0257 | -55.8927 | 2690 | 0.0000 | -1.4847 | -1.3840 | - |
| 'Intercept descendent model' | -1.4250 | 0.0279 | -51.1539 | 2690 | 0.0000 | -1.4796 | -1.3704 | - |
| 'Wake vs non-UBE' | 0.0094 | 0.0379 | 0.2471 | 2690 | 0.8048 | -0.0649 | 0.0836 | 0.8522 |
| 'UBE vs non-UBE' | 0.0578 | 0.0601 | 0.9624 | 2690 | 0.3359 | -0.0600 | 0.1755 | 0.6718 |
| 'UBE vs Wake' | 0.0484 | 0.0641 | 0.7559 | 2690 | 0.4498 | -0.0772 | 0.1741 | 0.7360 |
| **THETA BAND** | | | | | | | | |
| 'Intercept ascendent model' | -2.5583 | 0.0404 | -63.2505 | 2690 | 0.0000 | -2.6376 | -2.4789 | - |
| 'Intercept descendent model' | -2.6356 | 0.0440 | -59.9243 | 2690 | 0.0000 | -2.7218 | -2.5493 | - |
| 'Wake vs non-UBE' | -0.0773 | 0.0598 | -1.2941 | 2690 | 0.1957 | -0.1945 | 0.0398 | 0.4543 |
| 'UBE vs non-UBE' | -0.1135 | 0.0766 | -1.4820 | 2690 | 0.1384 | -0.2637 | 0.0367 | 0.4543 |
| 'UBE vs Wake' | -0.0362 | 0.0885 | -0.4092 | 2690 | 0.6824 | -0.2097 | 0.1373 | 0.8522 |
| **ALPHA BAND** | | | | | | | | |
| 'Intercept ascendent model' | -2.1277 | 0.0713 | -29.8616 | 2690 | 0.0000 | -2.2674 | -1.9880 | - |
| 'Intercept descendent model' | -1.9386 | 0.0805 | -24.0804 | 2690 | 0.0000 | -2.0964 | -1.7807 | - |
| 'Wake vs non-UBE' | 0.1891 | 0.1075 | 1.7593 | 2690 | 0.0786 | -0.0217 | 0.3999 | 0.3539 |
| 'UBE vs non-UBE' | -0.1591 | 0.0819 | -1.9426 | 2690 | 0.0522 | -0.3197 | 0.0015 | 0.3130 |
| 'UBE vs Wake' | -0.3482 | 0.1245 | -2.7974 | 2690 | 0.0052 | -0.5924 | -0.1041 | 0.0467 |
| **SIGMA BAND** | | | | | | | | |
| 'Intercept ascendent model' | -1.5461 | 0.0319 | -48.4574 | 2690 | 0.0000 | -1.6087 | -1.4835 | - |
| 'Intercept descendent model' | -1.5685 | 0.0348 | -45.1004 | 2690 | 0.0000 | -1.6367 | -1.5003 | - |
| 'Wake vs non-UBE' | -0.0224 | 0.0472 | -0.4749 | 2690 | 0.6349 | -0.1150 | 0.0701 | 0.8522 |
| 'UBE vs non-UBE' | -0.0480 | 0.0628 | -0.7644 | 2690 | 0.4447 | -0.1713 | 0.0752 | 0.7360 |
| 'UBE vs Wake' | -0.0256 | 0.0709 | -0.3611 | 2690 | 0.7180 | -0.1647 | 0.1135 | 0.8522 |
| **BETA BAND** | | | | | | | | |
| 'Intercept ascendent model' | -4.9350 | 0.0584 | -84.4583 | 2690 | 0.0000 | -5.0495 | -4.8204 | - |
| 'Intercept descendent model' | -4.9578 | 0.0668 | -74.2542 | 2690 | 0.0000 | -5.0887 | -4.8269 | - |
| 'Wake vs non-UBE' | -0.0228 | 0.0887 | -0.2575 | 2690 | 0.7968 | -0.1968 | 0.1511 | 0.8522 |
| 'UBE vs non-UBE' | -0.0264 | 0.0558 | -0.4732 | 2690 | 0.6361 | -0.1359 | 0.0831 | 0.8522 |
| 'UBE vs Wake' | -0.0036 | 0.0981 | -0.0364 | 2690 | 0.9709 | -0.1959 | 0.1887 | 0.9709 |
| **GAMMA BAND** | | | | | | | | |
| 'Intercept ascendent model' | -4.6887 | 0.0460 | -102.0224 | 2690 | 0.0000 | -4.7788 | -4.5986 | - |

| | | | | | | | | |
|---|---|---|---|---|---|---|---|---|
| **'Intercept descendent model'** | -4.5979 | 0.0532 | -86.3529 | 2690 | 0.0000 | -4.7023 | -4.4935 | - |
| **'Wake vs non-UBE'** | 0.0908 | 0.0703 | 1.2910 | 2690 | 0.1968 | -0.0471 | 0.2287 | 0.4543 |
| **'UBE vs non-UBE'** | 0.1865 | 0.0356 | 5.2361 | 2690 | 0.0000 | 0.1166 | 0.2563 | 0.0000 |
| **'UBE vs Wake'** | 0.0956 | 0.0749 | 1.2764 | 2690 | 0.2019 | -0.0513 | 0.2426 | 0.4543 |

### Hierarchy – Broadband comparisons

##### Table S20: Broadband analysis – full UBE sample (T=10% of max autocorrelation)

| | | | | | | **CI 95%** | | |
|---|---|---|---|---|---|---|---|---|
| **Contrast** | **Estimate** | **SE** | **tStat** | **DF** | **pvalue** | **lower** | **upper** | **FDR-adjusted p** |
| **'Intercept ascendent model'** | -5.5094 | 0.1233 | -44.6810 | 2690 | 0.0000 | -5.7512 | -5.2676 | - |
| **'Intercept descendent model'** | -5.5796 | 0.1436 | -38.8536 | 2690 | 0.0000 | -5.8611 | -5.2980 | - |
| **'Wake vs non-UBE'** | -0.0702 | 0.1893 | -0.3706 | 2690 | 0.7109 | -0.4413 | 0.3010 | 0.7109 |
| **'UBE vs non-UBE'** | -0.3624 | 0.0913 | -3.9683 | 2690 | 0.0001 | -0.5415 | -0.1833 | 0.0002 |
| **'UBE vs Wake'** | -0.2923 | 0.2003 | -1.4595 | 2690 | 0.1445 | -0.6850 | 0.1004 | 0.2168 |

##### Table S21: Broadband analysis – out-of-body experiences (T=10% of max autocorrelation)

| | | | | | | **CI 95%** | | |
|---|---|---|---|---|---|---|---|---|
| **Contrast** | **Estimate** | **SE** | **tStat** | **DF** | **pvalue** | **lower** | **upper** | **FDR-adjusted p** |
| **'Intercept ascendent model'** | -5.6598 | 0.1763 | -32.1059 | 870 | 0.0000 | -6.0058 | -5.3138 | - |
| **'Intercept descendent model'** | -5.4423 | 0.1927 | -28.2462 | 870 | 0.0000 | -5.8204 | -5.0641 | - |
| **'Wake vs non-UBE'** | 0.2176 | 0.2611 | 0.8331 | 870 | 0.4050 | -0.2950 | 0.7301 | 0.4050 |
| **'UBE vs non-UBE'** | -0.5752 | 0.1618 | -3.5540 | 870 | 0.0004 | -0.8928 | -0.2575 | 0.0012 |
| **'UBE vs Wake'** | -0.7928 | 0.2840 | -2.7912 | 870 | 0.0054 | -1.3502 | -0.2353 | 0.0080 |

### Hierarchy – Broadband sensitivity analyses

##### Table S22: Broadband sensitivity analysis – full UBE sample (T=0% of max autocorrelation)

| | | | | | | **CI 95%** | | |
|---|---|---|---|---|---|---|---|---|
| **Contrast** | **Estimate** | **SE** | **tStat** | **DF** | **pvalue** | **lower** | **upper** | **FDR-adjusted p** |
| **'Intercept ascendent model'** | -5.5602 | 0.1127 | -49.3481 | 2690 | 0.0000 | -5.7812 | -5.3393 | - |
| **'Intercept descendent model'** | -5.5666 | 0.1304 | -42.6942 | 2690 | 0.0000 | -5.8222 | -5.3109 | - |
| **'Wake vs non-UBE'** | -0.0063 | 0.1723 | -0.0365 | 2690 | 0.9709 | -0.3442 | 0.3316 | 0.9709 |
| **'UBE vs non-UBE'** | -0.3658 | 0.0885 | -4.1325 | 2690 | 0.0000 | -0.5394 | -0.1922 | 0.0001 |
| **'UBE vs Wake'** | -0.3595 | 0.1840 | -1.9538 | 2690 | 0.0508 | -0.7203 | 0.0013 | 0.0762 |

##### Table S23: Broadband sensitivity analysis – full UBE sample (T=20% of max autocorrelation)

| | | | | | | **CI 95%** |
|---|---|---|---|---|---|---|

| Contrast | Estimate | SE | tStat | DF | pvalue | lower | upper | FDR-adjusted p |
|---|---|---|---|---|---|---|---|---|
| 'Intercept ascendent model' | -5.5338 | 0.1171 | -47.2517 | 2690 | 0.0000 | -5.7635 | -5.3042 | - |
| 'Intercept descendent model' | -5.6235 | 0.1356 | -41.4599 | 2690 | 0.0000 | -5.8895 | -5.3575 | - |
| 'Wake vs non-UBE' | -0.0896 | 0.1792 | -0.5002 | 2690 | 0.6169 | -0.4410 | 0.2617 | 0.6169 |
| 'UBE vs non-UBE' | -0.3822 | 0.0869 | -4.3978 | 2690 | 0.0000 | -0.5526 | -0.2118 | 0.0000 |
| 'UBE vs Wake' | -0.2926 | 0.1900 | -1.5397 | 2690 | 0.1237 | -0.6652 | 0.0800 | 0.1856 |

**Table S24: Broadband sensitivity analysis – full UBE sample (T=30% of max autocorrelation)**

| | | | | | | CI 95% | | |
|---|---|---|---|---|---|---|---|---|
| Contrast | Estimate | SE | tStat | DF | pvalue | lower | upper | FDR-adjusted p |
| 'Intercept ascendent model' | -5.7109 | 0.1142 | -49.9887 | 2690 | 0.0000 | -5.9350 | -5.4869 | - |
| 'Intercept descendent model' | -5.8211 | 0.1320 | -44.0865 | 2690 | 0.0000 | -6.0800 | -5.5622 | - |
| 'Wake vs non-UBE' | -0.1101 | 0.1746 | -0.6307 | 2690 | 0.5283 | -0.4525 | 0.2323 | 0.5283 |
| 'UBE vs non-UBE' | -0.3523 | 0.0840 | -4.1918 | 2690 | 0.0000 | -0.5171 | -0.1875 | 0.0001 |
| 'UBE vs Wake' | -0.2422 | 0.1851 | -1.3080 | 2690 | 0.1910 | -0.6052 | 0.1209 | 0.2865 |

**Table S25: Broadband sensitivity analysis – full UBE sample (T=40% of max autocorrelation)**

| | | | | | | CI 95% | | |
|---|---|---|---|---|---|---|---|---|
| Contrast | Estimate | SE | tStat | DF | pvalue | lower | upper | FDR-adjusted p |
| 'Intercept ascendent model' | -5.7108 | 0.1131 | -50.4807 | 2690 | 0.0000 | -5.9327 | -5.4890 | - |
| 'Intercept descendent model' | -5.8611 | 0.1312 | -44.6754 | 2690 | 0.0000 | -6.1184 | -5.6039 | - |
| 'Wake vs non-UBE' | -0.1503 | 0.1732 | -0.8676 | 2690 | 0.3857 | -0.4900 | 0.1894 | 0.3857 |
| 'UBE vs non-UBE' | -0.4242 | 0.0841 | -5.0433 | 2690 | 0.0000 | -0.5892 | -0.2593 | 0.0000 |
| 'UBE vs Wake' | -0.2739 | 0.1837 | -1.4916 | 2690 | 0.1359 | -0.6341 | 0.0862 | 0.2039 |

**Table S26: Broadband sensitivity analysis – full UBE sample (T=50% of max autocorrelation)**

| | | | | | | CI 95% | | |
|---|---|---|---|---|---|---|---|---|
| Contrast | Estimate | SE | tStat | DF | pvalue | lower | upper | FDR-adjusted p |
| 'Intercept ascendent model' | -5.6839 | 0.1323 | -42.9735 | 2690 | 0.0000 | -5.9432 | -5.4245 | - |
| 'Intercept descendent model' | -5.4883 | 0.1547 | -35.4751 | 2690 | 0.0000 | -5.7917 | -5.1850 | - |
| 'Wake vs non-UBE' | 0.1955 | 0.2035 | 0.9607 | 2690 | 0.3368 | -0.2036 | 0.5946 | 0.3368 |
| 'UBE vs non-UBE' | -0.4128 | 0.0882 | -4.6802 | 2690 | 0.0000 | -0.5858 | -0.2399 | 0.0000 |
| 'UBE vs Wake' | -0.6084 | 0.2130 | -2.8562 | 2690 | 0.0043 | -1.0260 | -0.1907 | 0.0065 |

**Table S27: Broadband sensitivity analysis – full UBE sample (T=10% of max autocorrelation, using only 20% highest irreversibility values)**

| | | | | | | CI 95% | | |
|---|---|---|---|---|---|---|---|---|
| Contrast | Estimate | SE | tStat | DF | pvalue | lower | upper | FDR-adjusted p |
| 'Intercept ascendent model' | -5.3497 | 0.1272 | -42.0528 | 2690 | 0.0000 | -5.5992 | -5.1003 | - |
| 'Intercept descendent model' | -5.4244 | 0.1464 | -37.0400 | 2690 | 0.0000 | -5.7115 | -5.1372 | - |
| 'Wake vs non-UBE' | -0.0747 | 0.1940 | -0.3849 | 2690 | 0.7004 | -0.4550 | 0.3057 | 0.7004 |

| | | | | | | | | |
|---|---|---|---|---|---|---|---|---|
| 'UBE vs non-UBE' | -0.2875 | 0.1115 | -2.5786 | 2690 | 0.0100 | -0.5061 | -0.0689 | 0.0299 |
| 'UBE vs Wake' | -0.2128 | 0.2106 | -1.0109 | 2690 | 0.3122 | -0.6257 | 0.2000 | 0.4682 |

**Table S28: Broadband sensitivity analysis – full UBE sample (T=50% of max autocorrelation, using only 20% highest irreversibility values)**

| | | | | | | CI 95% | | |
|---|---|---|---|---|---|---|---|---|
| **Contrast** | **Estimate** | **SE** | **tStat** | **DF** | **pvalue** | **lower** | **upper** | **FDR-adjusted p** |
| 'Intercept ascendent model' | -5.4878 | 0.1348 | -40.7167 | 2690 | 0.0000 | -5.7521 | -5.2235 | - |
| 'Intercept descendent model' | -5.3066 | 0.1551 | -34.2129 | 2690 | 0.0000 | -5.6107 | -5.0024 | - |
| 'Wake vs non-UBE' | 0.1812 | 0.2055 | 0.8820 | 2690 | 0.3779 | -0.2217 | 0.5842 | 0.3779 |
| 'UBE vs non-UBE' | -0.3794 | 0.1076 | -3.5255 | 2690 | 0.0004 | -0.5905 | -0.1684 | 0.0013 |
| 'UBE vs Wake' | -0.5607 | 0.2204 | -2.5437 | 2690 | 0.0110 | -0.9929 | -0.1285 | 0.0165 |

**Table S29: Broadband sensitivity analysis – REM sleep group (T=10% of max autocorrelation)**

| | | | | | | CI 95% | | |
|---|---|---|---|---|---|---|---|---|
| **Contrast** | **Estimate** | **SE** | **tStat** | **DF** | **pvalue** | **lower** | **upper** | **FDR-adjusted p** |
| 'Intercept ascendent model' | -5.4994 | 0.1789 | -30.7393 | 605 | 0.0000 | -5.8508 | -5.1481 | - |
| 'Intercept descendent model' | -5.3264 | 0.1798 | -29.6251 | 605 | 0.0000 | -5.6795 | -4.9733 | - |
| 'Wake vs non-UBE' | 0.1730 | 0.0978 | 1.7693 | 605 | 0.0774 | -0.0190 | 0.3650 | 0.0774 |
| 'UBE vs non-UBE' | -0.4549 | 0.2226 | -2.0438 | 605 | 0.0414 | -0.8920 | -0.0178 | 0.0621 |
| 'UBE vs Wake' | -0.6278 | 0.2211 | -2.8391 | 605 | 0.0047 | -1.0622 | -0.1935 | 0.0140 |

**Table S30: Broadband sensitivity analysis – Light sleep group (T=10% of max autocorrelation)**

| | | | | | | CI 95% | | |
|---|---|---|---|---|---|---|---|---|
| **Contrast** | **Estimate** | **SE** | **tStat** | **DF** | **pvalue** | **lower** | **upper** | **FDR-adjusted p** |
| 'Intercept ascendent model' | -5.0697 | 0.0668 | -75.8415 | 625 | 0.0000 | -5.2009 | -4.9384 | - |
| 'Intercept descendent model' | -6.0134 | 0.0676 | -88.9034 | 625 | 0.0000 | -6.1462 | -5.8806 | - |
| 'Wake vs non-UBE' | -0.9437 | 0.0951 | -9.9236 | 625 | 0.0000 | -1.1305 | -0.7570 | 0.0000 |
| 'UBE vs non-UBE' | -0.4230 | 0.2068 | -2.0456 | 625 | 0.0412 | -0.8291 | -0.0169 | 0.0412 |
| 'UBE vs Wake' | 0.5207 | 0.2071 | 2.5146 | 625 | 0.0122 | 0.1140 | 0.9273 | 0.0183 |

**Table S31: Broadband sensitivity analysis – Meditation group (T=10% of max autocorrelation)**

| | | | | | | CI 95% | | |
|---|---|---|---|---|---|---|---|---|
| **Contrast** | **Estimate** | **SE** | **tStat** | **DF** | **pvalue** | **lower** | **upper** | **FDR-adjusted p** |
| 'Intercept ascendent model' | -5.8437 | 0.1615 | -36.1784 | 873 | 0.0000 | -6.1607 | -5.5267 | - |
| 'Intercept descendent model' | -5.7246 | 0.1617 | -35.4020 | 873 | 0.0000 | -6.0420 | -5.4073 | - |
| 'Wake vs non-UBE' | 0.1191 | 0.0731 | 1.6292 | 873 | 0.1036 | -0.0244 | 0.2625 | 0.1036 |
| 'UBE vs non-UBE' | -0.5675 | 0.1365 | -4.1583 | 873 | 0.0000 | -0.8353 | -0.2996 | 0.0001 |
| 'UBE vs Wake' | -0.6865 | 0.1361 | -5.0427 | 873 | 0.0000 | -0.9537 | -0.4193 | 0.0000 |

**Table S32: Broadband sensitivity analysis – Sleep arousal group (T=10% of max autocorrelation)**

| | | | | | | CI 95% | | |
|---|---|---|---|---|---|---|---|---|
| **Contrast** | **Estimate** | **SE** | **tStat** | **DF** | **pvalue** | **lower** | **upper** | **FDR-adjusted p** |
| **'Intercept ascendent model'** | -5.2165 | 0.1767 | -29.5293 | 864 | 0.0000 | -5.5632 | -4.8698 | - |
| **'Intercept descendent model'** | -5.6923 | 0.1770 | -32.1533 | 864 | 0.0000 | -6.0397 | -5.3448 | - |
| **'Wake vs non-UBE'** | -0.4757 | 0.0816 | -5.8280 | 864 | 0.0000 | -0.6359 | -0.3155 | 0.0000 |
| **'UBE vs non-UBE'** | -0.4962 | 0.1713 | -2.8964 | 864 | 0.0039 | -0.8325 | -0.1600 | 0.0058 |
| **'UBE vs Wake'** | -0.0205 | 0.1656 | -0.1239 | 864 | 0.9014 | -0.3455 | 0.3045 | 0.9014 |

### Hierarchy – Frequency band comparisons

**Table S33: Frequency band analysis – full UBE sample (T=10% of max autocorrelation)**

| | | | | | | CI 95% | | |
|---|---|---|---|---|---|---|---|---|
| **Contrast** | **Estimate** | **SE** | **tStat** | **DF** | **pvalue** | **lower** | **upper** | **FDR-adjusted p** |
| **DELTA BAND** | | | | | | | | |
| **'Intercept ascendent model'** | -2.4068 | 0.0310 | -77.5254 | 2690 | 0.0000 | -2.4677 | -2.3460 | - |
| **'Intercept descendent model'** | -2.3444 | 0.0337 | -69.5460 | 2690 | 0.0000 | -2.4105 | -2.2783 | - |
| **'Wake vs non-UBE'** | 0.0624 | 0.0458 | 1.3619 | 2690 | 0.1733 | -0.0274 | 0.1523 | 0.4711 |
| **'UBE vs non-UBE'** | 0.0510 | 0.0704 | 0.7252 | 2690 | 0.4684 | -0.0869 | 0.1890 | 0.6643 |
| **'UBE vs Wake'** | -0.0114 | 0.0758 | -0.1502 | 2690 | 0.8806 | -0.1601 | 0.1373 | 0.8806 |
| **THETA BAND** | | | | | | | | |
| **'Intercept ascendent model'** | -4.2826 | 0.0528 | -81.1369 | 2690 | 0.0000 | -4.3861 | -4.1791 | - |
| **'Intercept descendent model'** | -4.4269 | 0.0576 | -76.8827 | 2690 | 0.0000 | -4.5398 | -4.3140 | - |
| **'Wake vs non-UBE'** | -0.1444 | 0.0781 | -1.8481 | 2690 | 0.0647 | -0.2975 | 0.0088 | 0.2329 |
| **'UBE vs non-UBE'** | -0.1089 | 0.0918 | -1.1865 | 2690 | 0.2355 | -0.2888 | 0.0711 | 0.4711 |
| **'UBE vs Wake'** | 0.0355 | 0.1099 | 0.3227 | 2690 | 0.7470 | -0.1801 | 0.2510 | 0.7909 |
| **ALPHA BAND** | | | | | | | | |
| **'Intercept ascendent model'** | -4.6115 | 0.0959 | -48.0966 | 2690 | 0.0000 | -4.7995 | -4.4235 | - |
| **'Intercept descendent model'** | -4.8024 | 0.1090 | -44.0457 | 2690 | 0.0000 | -5.0162 | -4.5886 | - |
| **'Wake vs non-UBE'** | -0.1909 | 0.1452 | -1.3148 | 2690 | 0.1887 | -0.4756 | 0.0938 | 0.4711 |
| **'UBE vs non-UBE'** | -0.2823 | 0.1049 | -2.6924 | 2690 | 0.0071 | -0.4879 | -0.0767 | 0.0573 |
| **'UBE vs Wake'** | -0.0914 | 0.1653 | -0.5531 | 2690 | 0.5802 | -0.4155 | 0.2327 | 0.6770 |
| **SIGMA BAND** | | | | | | | | |
| **'Intercept ascendent model'** | -5.0812 | 0.0650 | -78.1492 | 2690 | 0.0000 | -5.2087 | -4.9537 | - |
| **'Intercept descendent model'** | -4.8766 | 0.0720 | -67.7541 | 2690 | 0.0000 | -5.0177 | -4.7354 | - |
| **'Wake vs non-UBE'** | 0.2046 | 0.0970 | 2.1093 | 2690 | 0.0350 | 0.0144 | 0.3948 | 0.1576 |
| **'UBE vs non-UBE'** | -0.1107 | 0.0900 | -1.2296 | 2690 | 0.2190 | -0.2871 | 0.0658 | 0.4711 |
| **'UBE vs Wake'** | -0.3153 | 0.1216 | -2.5937 | 2690 | 0.0095 | -0.5536 | -0.0769 | 0.0573 |
| **BETA BAND** | | | | | | | | |
| **'Intercept ascendent model'** | -5.9711 | 0.0798 | -74.7834 | 2690 | 0.0000 | -6.1277 | -5.8146 | - |
| **'Intercept descendent model'** | -6.0878 | 0.0922 | -66.0115 | 2690 | 0.0000 | -6.2686 | -5.9070 | - |
| **'Wake vs non-UBE'** | -0.1167 | 0.1220 | -0.9565 | 2690 | 0.3389 | -0.3559 | 0.1225 | 0.5546 |

| | | | | | | | | |
|---|---|---|---|---|---|---|---|---|
| **'UBE vs non-UBE'** | -0.1989 | 0.0668 | -2.9789 | 2690 | 0.0029 | -0.3298 | -0.0680 | 0.0525 |
| **'UBE vs Wake'** | -0.0822 | 0.1314 | -0.6260 | 2690 | 0.5314 | -0.3398 | 0.1753 | 0.6770 |
| **GAMMA BAND** | | | | | | | | |
| **'Intercept ascendent model'** | -6.2175 | 0.0470 | -132.3751 | 2690 | 0.0000 | -6.3096 | -6.1254 | - |
| **'Intercept descendent model'** | -6.2877 | 0.0539 | -116.7556 | 2690 | 0.0000 | -6.3933 | -6.1821 | - |
| **'Wake vs non-UBE'** | -0.0702 | 0.0715 | -0.9830 | 2690 | 0.3257 | -0.2104 | 0.0699 | 0.5546 |
| **'UBE vs non-UBE'** | -0.0296 | 0.0419 | -0.7067 | 2690 | 0.4798 | -0.1116 | 0.0525 | 0.6643 |
| **'UBE vs Wake'** | 0.0407 | 0.0779 | 0.5219 | 2690 | 0.6018 | -0.1121 | 0.1935 | 0.6770 |

**Table S34: Frequency band analysis – out-of-body experiences (T=10% of max autocorrelation)**

| | | | | | | **CI 95%** | | |
|---|---|---|---|---|---|---|---|---|
| **Contrast** | **Estimate** | **SE** | **tStat** | **DF** | **pvalue** | **lower** | **upper** | **FDR-adjusted p** |
| **DELTA BAND** | | | | | | | | |
| **'Intercept ascendent model'** | -2.4184 | 0.0550 | -43.9902 | 870 | 0.0000 | -2.5263 | -2.3105 | - |
| **'Intercept descendent model'** | -2.5507 | 0.0558 | -45.7181 | 870 | 0.0000 | -2.6602 | -2.4412 | - |
| **'Wake vs non-UBE'** | -0.1323 | 0.0783 | -1.6889 | 870 | 0.0916 | -0.2860 | 0.0214 | 0.3298 |
| **'UBE vs non-UBE'** | -0.1514 | 0.1256 | -1.2054 | 870 | 0.2284 | -0.3978 | 0.0951 | 0.5138 |
| **'UBE vs Wake'** | -0.0191 | 0.1334 | -0.1430 | 870 | 0.8863 | -0.2809 | 0.2427 | 0.9517 |
| **THETA BAND** | | | | | | | | |
| **'Intercept ascendent model'** | -4.1135 | 0.0671 | -61.2949 | 870 | 0.0000 | -4.2452 | -3.9818 | - |
| **'Intercept descendent model'** | -4.2821 | 0.0682 | -62.7559 | 870 | 0.0000 | -4.4160 | -4.1482 | - |
| **'Wake vs non-UBE'** | -0.1686 | 0.0957 | -1.7615 | 870 | 0.0785 | -0.3564 | 0.0193 | 0.3298 |
| **'UBE vs non-UBE'** | -0.1835 | 0.1436 | -1.2779 | 870 | 0.2016 | -0.4653 | 0.0983 | 0.5138 |
| **'UBE vs Wake'** | -0.0149 | 0.1564 | -0.0953 | 870 | 0.9241 | -0.3220 | 0.2922 | 0.9517 |
| **ALPHA BAND** | | | | | | | | |
| **'Intercept ascendent model'** | -4.5511 | 0.1580 | -28.8103 | 870 | 0.0000 | -4.8611 | -4.2411 | - |
| **'Intercept descendent model'** | -4.8813 | 0.1683 | -29.0046 | 870 | 0.0000 | -5.2117 | -4.5510 | - |
| **'Wake vs non-UBE'** | -0.3303 | 0.2308 | -1.4308 | 870 | 0.1528 | -0.7833 | 0.1228 | 0.4585 |
| **'UBE vs non-UBE'** | -0.5144 | 0.2006 | -2.5645 | 870 | 0.0105 | -0.9081 | -0.1207 | 0.1890 |
| **'UBE vs Wake'** | -0.1842 | 0.2729 | -0.6747 | 870 | 0.5000 | -0.7199 | 0.3515 | 0.7250 |
| **SIGMA BAND** | | | | | | | | |
| **'Intercept ascendent model'** | -4.9747 | 0.1217 | -40.8680 | 870 | 0.0000 | -5.2136 | -4.7358 | - |
| **'Intercept descendent model'** | -4.6537 | 0.1270 | -36.6537 | 870 | 0.0000 | -4.9029 | -4.4045 | - |
| **'Wake vs non-UBE'** | 0.3210 | 0.1759 | 1.8249 | 870 | 0.0684 | -0.0242 | 0.6662 | 0.3298 |
| **'UBE vs non-UBE'** | -0.1234 | 0.1713 | -0.7206 | 870 | 0.4713 | -0.4595 | 0.2127 | 0.7250 |
| **'UBE vs Wake'** | -0.4444 | 0.2209 | -2.0118 | 870 | 0.0445 | -0.8779 | -0.0109 | 0.3298 |
| **BETA BAND** | | | | | | | | |
| **'Intercept ascendent model'** | -6.6098 | 0.1435 | -46.0667 | 870 | 0.0000 | -6.8914 | -6.3282 | - |
| **'Intercept descendent model'** | -6.6808 | 0.1557 | -42.9104 | 870 | 0.0000 | -6.9863 | -6.3752 | - |
| **'Wake vs non-UBE'** | -0.0709 | 0.2117 | -0.3351 | 870 | 0.7376 | -0.4865 | 0.3446 | 0.8852 |
| **'UBE vs non-UBE'** | -0.0850 | 0.1332 | -0.6381 | 870 | 0.5236 | -0.3464 | 0.1764 | 0.7250 |
| **'UBE vs Wake'** | -0.0140 | 0.2314 | -0.0606 | 870 | 0.9517 | -0.4683 | 0.4402 | 0.9517 |

| | | | | | | | | |
|---|---|---|---|---|---|---|---|---|
| | | | | **GAMMA BAND** | | | | |
| **'Intercept ascendent model'** | -6.0876 | 0.1117 | -54.5100 | 870 | 0.0000 | -6.3068 | -5.8684 | - |
| **'Intercept descendent model'** | -6.2246 | 0.1235 | -50.4050 | 870 | 0.0000 | -6.4670 | -5.9822 | - |
| **'Wake vs non-UBE'** | -0.1370 | 0.1665 | -0.8226 | 870 | 0.4110 | -0.4637 | 0.1898 | 0.7250 |
| **'UBE vs non-UBE'** | 0.0527 | 0.0914 | 0.5764 | 870 | 0.5645 | -0.1267 | 0.2320 | 0.7257 |
| **'UBE vs Wake'** | 0.1896 | 0.1773 | 1.0696 | 870 | 0.2851 | -0.1583 | 0.5376 | 0.5702 |

### Hierarchy – Frequency band sensitivity analyses

**Table S35: Frequency band sensitivity analysis – full UBE sample (T=50% of max autocorrelation)**

| | | | | | | **CI 95%** | | |
|---|---|---|---|---|---|---|---|---|
| **Contrast** | **Estimate** | **SE** | **tStat** | **DF** | **pvalue** | **lower** | **upper** | **FDR-adjusted p** |
| | | | | **DELTA BAND** | | | | |
| **'Intercept ascendent model'** | -1.9696 | 0.0282 | -69.8926 | 2690 | 0.0000 | -2.0249 | -1.9143 | - |
| **'Intercept descendent model'** | -1.9600 | 0.0306 | -64.0487 | 2690 | 0.0000 | -2.0200 | -1.9000 | - |
| **'Wake vs non-UBE'** | 0.0096 | 0.0416 | 0.2310 | 2690 | 0.8174 | -0.0720 | 0.0912 | 0.8930 |
| **'UBE vs non-UBE'** | 0.0837 | 0.0635 | 1.3183 | 2690 | 0.1875 | -0.0408 | 0.2081 | 0.5343 |
| **'UBE vs Wake'** | 0.0741 | 0.0685 | 1.0810 | 2690 | 0.2798 | -0.0603 | 0.2084 | 0.5343 |
| | | | | **THETA BAND** | | | | |
| **'Intercept ascendent model'** | -3.0497 | 0.0415 | -73.4402 | 2690 | 0.0000 | -3.1311 | -2.9683 | - |
| **'Intercept descendent model'** | -3.1231 | 0.0451 | -69.2023 | 2690 | 0.0000 | -3.2116 | -3.0346 | - |
| **'Wake vs non-UBE'** | -0.0734 | 0.0613 | -1.1975 | 2690 | 0.2312 | -0.1937 | 0.0468 | 0.5343 |
| **'UBE vs non-UBE'** | -0.0830 | 0.0801 | -1.0355 | 2690 | 0.3005 | -0.2401 | 0.0741 | 0.5343 |
| **'UBE vs Wake'** | -0.0095 | 0.0919 | -0.1038 | 2690 | 0.9173 | -0.1897 | 0.1706 | 0.9173 |
| | | | | **ALPHA BAND** | | | | |
| **'Intercept ascendent model'** | -2.6369 | 0.0685 | -38.5125 | 2690 | 0.0000 | -2.7712 | -2.5027 | - |
| **'Intercept descendent model'** | -2.4695 | 0.0769 | -32.1033 | 2690 | 0.0000 | -2.6204 | -2.3187 | - |
| **'Wake vs non-UBE'** | 0.1674 | 0.1030 | 1.6254 | 2690 | 0.1042 | -0.0345 | 0.3693 | 0.4688 |
| **'UBE vs non-UBE'** | -0.1399 | 0.0843 | -1.6590 | 2690 | 0.0972 | -0.3053 | 0.0255 | 0.4688 |
| **'UBE vs Wake'** | -0.3073 | 0.1220 | -2.5184 | 2690 | 0.0118 | -0.5466 | -0.0680 | 0.1066 |
| | | | | **SIGMA BAND** | | | | |
| **'Intercept ascendent model'** | -2.0765 | 0.0315 | -65.8663 | 2690 | 0.0000 | -2.1383 | -2.0147 | - |
| **'Intercept descendent model'** | -2.0857 | 0.0343 | -60.7909 | 2690 | 0.0000 | -2.1530 | -2.0184 | - |
| **'Wake vs non-UBE'** | -0.0092 | 0.0466 | -0.1976 | 2690 | 0.8434 | -0.1006 | 0.0822 | 0.8930 |
| **'UBE vs non-UBE'** | -0.0610 | 0.0661 | -0.9228 | 2690 | 0.3562 | -0.1907 | 0.0686 | 0.5343 |
| **'UBE vs Wake'** | -0.0518 | 0.0729 | -0.7103 | 2690 | 0.4776 | -0.1948 | 0.0912 | 0.6613 |
| | | | | **BETA BAND** | | | | |
| **'Intercept ascendent model'** | -5.2468 | 0.0584 | -89.8526 | 2690 | 0.0000 | -5.3613 | -5.1323 | - |
| **'Intercept descendent model'** | -5.2773 | 0.0663 | -79.6427 | 2690 | 0.0000 | -5.4072 | -5.1474 | - |
| **'Wake vs non-UBE'** | -0.0305 | 0.0883 | -0.3451 | 2690 | 0.7301 | -0.2037 | 0.1427 | 0.8930 |
| **'UBE vs non-UBE'** | -0.0583 | 0.0609 | -0.9567 | 2690 | 0.3388 | -0.1777 | 0.0612 | 0.5343 |

| | | | | | | | | |
|---|---|---|---|---|---|---|---|---|
| **'UBE vs Wake'** | -0.0278 | 0.0997 | -0.2787 | 2690 | 0.7805 | -0.2233 | 0.1677 | 0.8930 |
| | | | | **GAMMA BAND** | | | | |
| **'Intercept ascendent model'** | -4.8910 | 0.0431 | -113.4515 | 2690 | 0.0000 | -4.9755 | -4.8064 | - |
| **'Intercept descendent model'** | -4.8263 | 0.0494 | -97.7809 | 2690 | 0.0000 | -4.9230 | -4.7295 | - |
| **'Wake vs non-UBE'** | 0.0647 | 0.0655 | 0.9871 | 2690 | 0.3237 | -0.0638 | 0.1932 | 0.5343 |
| **'UBE vs non-UBE'** | 0.1547 | 0.0386 | 4.0098 | 2690 | 0.0001 | 0.0790 | 0.2303 | 0.0011 |
| **'UBE vs Wake'** | 0.0900 | 0.0716 | 1.2574 | 2690 | 0.2087 | -0.0504 | 0.2303 | 0.5343 |

**Table S36: Frequency band sensitivity analysis – full UBE sample (T=10% of max autocorrelation, using only 30% highest irreversibility values)**

| | | | | | | **CI 95%** | | |
|---|---|---|---|---|---|---|---|---|
| **Contrast** | **Estimate** | **SE** | **tStat** | **DF** | **pvalue** | **lower** | **upper** | **FDR-adjusted p** |
| | | | | **DELTA BAND** | | | | |
| **'Intercept ascendent model'** | -2.2280 | 0.0359 | -62.0793 | 2690 | 0.0000 | -2.2984 | -2.1577 | - |
| **'Intercept descendent model'** | -2.1797 | 0.0390 | -55.9517 | 2690 | 0.0000 | -2.2560 | -2.1033 | - |
| **'Wake vs non-UBE'** | 0.0484 | 0.0530 | 0.9135 | 2690 | 0.3611 | -0.0555 | 0.1522 | 0.5909 |
| **'UBE vs non-UBE'** | 0.0504 | 0.0807 | 0.6239 | 2690 | 0.5328 | -0.1079 | 0.2086 | 0.6626 |
| **'UBE vs Wake'** | 0.0020 | 0.0872 | 0.0226 | 2690 | 0.9820 | -0.1691 | 0.1730 | 0.9820 |
| | | | | **THETA BAND** | | | | |
| **'Intercept ascendent model'** | -4.1537 | 0.0592 | -70.1751 | 2690 | 0.0000 | -4.2698 | -4.0377 | - |
| **'Intercept descendent model'** | -4.3166 | 0.0645 | -66.9112 | 2690 | 0.0000 | -4.4431 | -4.1901 | - |
| **'Wake vs non-UBE'** | -0.1629 | 0.0876 | -1.8605 | 2690 | 0.0629 | -0.3346 | 0.0088 | 0.2265 |
| **'UBE vs non-UBE'** | -0.0885 | 0.1058 | -0.8371 | 2690 | 0.4026 | -0.2959 | 0.1189 | 0.6039 |
| **'UBE vs Wake'** | 0.0744 | 0.1250 | 0.5946 | 2690 | 0.5521 | -0.1708 | 0.3195 | 0.6626 |
| | | | | **ALPHA BAND** | | | | |
| **'Intercept ascendent model'** | -4.5103 | 0.0953 | -47.3440 | 2690 | 0.0000 | -4.6971 | -4.3235 | - |
| **'Intercept descendent model'** | -4.7112 | 0.1073 | -43.9174 | 2690 | 0.0000 | -4.9215 | -4.5008 | - |
| **'Wake vs non-UBE'** | -0.2009 | 0.1435 | -1.4004 | 2690 | 0.1615 | -0.4822 | 0.0804 | 0.4845 |
| **'UBE vs non-UBE'** | -0.2811 | 0.1179 | -2.3832 | 2690 | 0.0172 | -0.5123 | -0.0498 | 0.1034 |
| **'UBE vs Wake'** | -0.0801 | 0.1698 | -0.4720 | 2690 | 0.6370 | -0.4131 | 0.2528 | 0.7166 |
| | | | | **SIGMA BAND** | | | | |
| **'Intercept ascendent model'** | -4.8769 | 0.0671 | -72.6924 | 2690 | 0.0000 | -5.0084 | -4.7453 | - |
| **'Intercept descendent model'** | -4.6633 | 0.0738 | -63.1473 | 2690 | 0.0000 | -4.8081 | -4.5185 | - |
| **'Wake vs non-UBE'** | 0.2136 | 0.0998 | 2.1409 | 2690 | 0.0324 | 0.0180 | 0.4092 | 0.1457 |
| **'UBE vs non-UBE'** | -0.1023 | 0.0994 | -1.0296 | 2690 | 0.3033 | -0.2971 | 0.0925 | 0.5909 |
| **'UBE vs Wake'** | -0.3159 | 0.1293 | -2.4439 | 2690 | 0.0146 | -0.5694 | -0.0624 | 0.1034 |
| | | | | **BETA BAND** | | | | |
| **'Intercept ascendent model'** | -5.6331 | 0.0791 | -71.1925 | 2690 | 0.0000 | -5.7882 | -5.4779 | - |
| **'Intercept descendent model'** | -5.7501 | 0.0908 | -63.3597 | 2690 | 0.0000 | -5.9281 | -5.5722 | - |
| **'Wake vs non-UBE'** | -0.1170 | 0.1204 | -0.9719 | 2690 | 0.3312 | -0.3531 | 0.1191 | 0.5909 |
| **'UBE vs non-UBE'** | -0.2114 | 0.0722 | -2.9284 | 2690 | 0.0034 | -0.3530 | -0.0699 | 0.0618 |
| **'UBE vs Wake'** | -0.0944 | 0.1318 | -0.7164 | 2690 | 0.4738 | -0.3528 | 0.1640 | 0.6561 |

| | | | | | | | | |
|---|---|---|---|---|---|---|---|---|
| | | | | **GAMMA BAND** | | | | |
| **'Intercept ascendent model'** | -5.8172 | 0.0468 | -124.3334 | 2690 | 0.0000 | -5.9089 | -5.7254 | - |
| **'Intercept descendent model'** | -5.8966 | 0.0534 | -110.5258 | 2690 | 0.0000 | -6.0012 | -5.7919 | - |
| **'Wake vs non-UBE'** | -0.0794 | 0.0710 | -1.1189 | 2690 | 0.2633 | -0.2185 | 0.0597 | 0.5909 |
| **'UBE vs non-UBE'** | -0.0567 | 0.0445 | -1.2731 | 2690 | 0.2031 | -0.1441 | 0.0306 | 0.5222 |
| **'UBE vs Wake'** | 0.0227 | 0.0785 | 0.2890 | 2690 | 0.7726 | -0.1312 | 0.1766 | 0.8181 |

**Table S37: Frequency band sensitivity analysis – full UBE sample (T=50% of max autocorrelation, using only 30% highest irreversibility values)**

| | | | | | | **CI 95%** | | |
|---|---|---|---|---|---|---|---|---|
| **Contrast** | **Estimate** | **SE** | **tStat** | **DF** | **pvalue** | **lower** | **upper** | **FDR-adjusted p** |
| | | | | **DELTA BAND** | | | | |
| **'Intercept ascendent model'** | -1.7772 | 0.0340 | -52.2893 | 2690 | 0.0000 | -1.8438 | -1.7105 | - |
| **'Intercept descendent model'** | -1.7742 | 0.0369 | -48.0586 | 2690 | 0.0000 | -1.8466 | -1.7018 | - |
| **'Wake vs non-UBE'** | 0.0030 | 0.0502 | 0.0588 | 2690 | 0.9531 | -0.0954 | 0.1013 | 0.9874 |
| **'UBE vs non-UBE'** | 0.0998 | 0.0749 | 1.3318 | 2690 | 0.1830 | -0.0471 | 0.2467 | 0.5281 |
| **'UBE vs Wake'** | 0.0968 | 0.0815 | 1.1886 | 2690 | 0.2347 | -0.0629 | 0.2565 | 0.5281 |
| | | | | **THETA BAND** | | | | |
| **'Intercept ascendent model'** | -2.9135 | 0.0457 | -63.7776 | 2690 | 0.0000 | -3.0030 | -2.8239 | - |
| **'Intercept descendent model'** | -2.9869 | 0.0496 | -60.2381 | 2690 | 0.0000 | -3.0842 | -2.8897 | - |
| **'Wake vs non-UBE'** | -0.0735 | 0.0674 | -1.0898 | 2690 | 0.2759 | -0.2057 | 0.0587 | 0.5447 |
| **'UBE vs non-UBE'** | -0.0262 | 0.0934 | -0.2801 | 2690 | 0.7794 | -0.2093 | 0.1570 | 0.8768 |
| **'UBE vs Wake'** | 0.0473 | 0.1047 | 0.4517 | 2690 | 0.6515 | -0.1581 | 0.2527 | 0.8290 |
| | | | | **ALPHA BAND** | | | | |
| **'Intercept ascendent model'** | -2.5254 | 0.0688 | -36.7054 | 2690 | 0.0000 | -2.6603 | -2.3905 | - |
| **'Intercept descendent model'** | -2.3979 | 0.0765 | -31.3407 | 2690 | 0.0000 | -2.5479 | -2.2479 | - |
| **'Wake vs non-UBE'** | 0.1275 | 0.1029 | 1.2390 | 2690 | 0.2155 | -0.0743 | 0.3292 | 0.5281 |
| **'UBE vs non-UBE'** | -0.1247 | 0.0969 | -1.2868 | 2690 | 0.1983 | -0.3148 | 0.0653 | 0.5281 |
| **'UBE vs Wake'** | -0.2522 | 0.1286 | -1.9613 | 2690 | 0.0499 | -0.5044 | -0.0001 | 0.4495 |
| | | | | **SIGMA BAND** | | | | |
| **'Intercept ascendent model'** | -1.8846 | 0.0331 | -56.9046 | 2690 | 0.0000 | -1.9496 | -1.8197 | - |
| **'Intercept descendent model'** | -1.8839 | 0.0360 | -52.3373 | 2690 | 0.0000 | -1.9545 | -1.8133 | - |
| **'Wake vs non-UBE'** | 0.0008 | 0.0489 | 0.0157 | 2690 | 0.9874 | -0.0951 | 0.0967 | 0.9874 |
| **'UBE vs non-UBE'** | -0.0785 | 0.0764 | -1.0264 | 2690 | 0.3048 | -0.2283 | 0.0714 | 0.5447 |
| **'UBE vs Wake'** | -0.0792 | 0.0818 | -0.9685 | 2690 | 0.3329 | -0.2396 | 0.0812 | 0.5447 |
| | | | | **BETA BAND** | | | | |
| **'Intercept ascendent model'** | -4.8887 | 0.0583 | -83.8162 | 2690 | 0.0000 | -5.0031 | -4.7744 | - |
| **'Intercept descendent model'** | -4.9237 | 0.0657 | -74.8994 | 2690 | 0.0000 | -5.0526 | -4.7948 | - |
| **'Wake vs non-UBE'** | -0.0350 | 0.0879 | -0.3977 | 2690 | 0.6909 | -0.2073 | 0.1374 | 0.8290 |
| **'UBE vs non-UBE'** | -0.0791 | 0.0664 | -1.1916 | 2690 | 0.2335 | -0.2093 | 0.0511 | 0.5281 |
| **'UBE vs Wake'** | -0.0442 | 0.1017 | -0.4341 | 2690 | 0.6643 | -0.2437 | 0.1553 | 0.8290 |
| | | | | **GAMMA BAND** | | | | |

| | | | | | | | | |
|---|---|---|---|---|---|---|---|---|
| **'Intercept ascendent model'** | -4.4769 | 0.0421 | -106.3508 | 2690 | 0.0000 | -4.5594 | -4.3943 | - |
| **'Intercept descendent model'** | -4.4260 | 0.0478 | -92.5358 | 2690 | 0.0000 | -4.5198 | -4.3323 | - |
| **'Wake vs non-UBE'** | 0.0508 | 0.0637 | 0.7978 | 2690 | 0.4250 | -0.0741 | 0.1758 | 0.6376 |
| **'UBE vs non-UBE'** | 0.1373 | 0.0412 | 3.3351 | 2690 | 0.0009 | 0.0566 | 0.2180 | 0.0156 |
| **'UBE vs Wake'** | 0.0865 | 0.0710 | 1.2182 | 2690 | 0.2233 | -0.0527 | 0.2256 | 0.5281 |

### Complexity:

##### Table S38: full UBE sample

| | | | | | | **CI 95%** | | |
|---|---|---|---|---|---|---|---|---|
| **Contrast** | **Estimate** | **SE** | **tStat** | **DF** | **pvalue** | **lower** | **upper** | **FDR-adjusted p** |
| **'Intercept ascendent model'** | 0.2968 | 0.0058 | 50.7393 | 2690 | 0.0000 | 0.2853 | 0.3083 | - |
| **'Intercept descendent model'** | 0.3312 | 0.0070 | 47.1969 | 2690 | 0.0000 | 0.3174 | 0.3450 | - |
| **'Wake vs non-UBE'** | 0.0344 | 0.0091 | 3.7656 | 2690 | 0.0002 | 0.0165 | 0.0523 | 0.0003 |
| **'UBE vs non-UBE'** | 0.0352 | 0.0026 | 13.7484 | 2690 | 0.0000 | 0.0302 | 0.0402 | 0.0000 |
| **'UBE vs Wake'** | 0.0008 | 0.0093 | 0.0824 | 2690 | 0.9343 | -0.0175 | 0.0190 | 0.9343 |

##### Table S39: out-of-body experiences

| | | | | | | **CI 95%** | | |
|---|---|---|---|---|---|---|---|---|
| **Contrast** | **Estimate** | **SE** | **tStat** | **DF** | **pvalue** | **lower** | **upper** | **FDR-adjusted p** |
| **'Intercept ascendent model'** | 0.3101 | 0.0077 | 40.0612 | 870 | 0.0000 | 0.2949 | 0.3253 | - |
| **'Intercept descendent model'** | 0.3259 | 0.0089 | 36.7873 | 870 | 0.0000 | 0.3085 | 0.3433 | - |
| **'Wake vs non-UBE'** | 0.0158 | 0.0118 | 1.3389 | 870 | 0.1810 | -0.0073 | 0.0388 | 0.2714 |
| **'UBE vs non-UBE'** | 0.0232 | 0.0045 | 5.1563 | 870 | 0.0000 | 0.0143 | 0.0320 | 0.0000 |
| **'UBE vs Wake'** | 0.0074 | 0.0121 | 0.6139 | 870 | 0.5394 | -0.0163 | 0.0311 | 0.5394 |
